# The MARK2 kinase acts as a gatekeeper of CD28-dependent co-stimulation in T cells

**DOI:** 10.1101/2025.10.13.682047

**Authors:** Hermine Ferran, Federico Marconi, Stephanie Dogniaux, Christel Goudot, Aymeric Zellner, Theo Cools, Thibaut Larcher, Elise Brisebard, Noémie Paillon, Frederic Fiore, CIPHE-GEMTISortium investigators, Claire Hivroz, Laurence Bataille

## Abstract

Naïve T cell activation requires not only antigen recognition through the TCR but also a co-stimulatory signal, mainly provided by CD28. Here, using a T cell–specific conditional knockout (cKO) model, we identify the microtubule-affinity kinase 2 (MARK2) as a key intracellular checkpoint that limits CD28-mediated co-stimulation. *In vivo*, MARK2 deficiency promotes the development of central memory T cells, enhances basal glycolysis activity in naïve CD8 T cells, and leads to the development of systemic autoimmunity in aged mice. In MARK2-deficient CD8 T cells, TCR engagement alone drives sustained proliferation, cytokine production, and glycolysis, processes that normally require CD28 co-stimulation. Single-cell transcriptomic analysis reveals that MARK2 regulates the expression of genes involved in CD28 signaling and metabolic switch. We show that MARK2 restrains the PI3K–AKT–mTORC1 pathway by limiting CD28-driven transcriptional and metabolic programs. Mechanistically, we demonstrate that MARK2 phosphorylates CREB regulated transcription coactivator 2 (CRTC2) and suppresses CREB-mediated transcription and mTOR activation, whereas CD28 engagement lifts this inhibition. Together, our results redefine the role of CD28 that not only amplifies TCR signaling but also relieves a MARK2-dependent inhibitory signal. This work provides new insights into T cell activation, metabolism and immune tolerance with potential implications for immunotherapeutic strategies in cancer and autoimmunity.

## Introduction

T cell activation is initiated upon antigen recognition by the T cell receptor (TCR) which triggers a signaling cascade essential for naïve T cell priming and for the development of an adaptive immune response ^1^. Although the TCR chains lack intrinsic signaling domains, signal transduction is mediated by the associated CD3 subunits (CD3γ, CD3δ, CD3ε, and CD3ζ), which contain one or more immunoreceptor tyrosine-based activation motifs (ITAMs). Upon TCR engagement, these ITAMs are rapidly phosphorylated by Src family kinases, enabling the recruitment and activation of the kinase ZAP70 (zeta-chain-associated protein kinase 70). ZAP70 then phosphorylates the adaptor protein LAT (linker for activation of T cells) on multiple tyrosine residues, promoting the assembly of a multiprotein signaling complex (called the LAT-signalosome) which contains GRB2, GADS, SLP-76, and PLCγ1 and is essential for initiating downstream signaling ^2^. However, TCR engagement alone is not sufficient to fully activate T cells. Co-stimulatory receptors such as CD28, which is constitutively expressed on naïve T cells, are essential to amplify TCR signaling and promote sustained activation. Upon interaction with its ligands B7-1 (CD80) or B7-2 (CD86) on antigen-presenting cells, CD28 activates the PI3K– Akt–mTOR pathway, to facilitate glucose uptake and metabolic reprogramming necessary for T cell proliferation, differentiation, and acquisition of effector functions ^3^.

T cell responses are tightly regulated by a network of surface co-receptors and intracellular regulators that control the strength and duration of the immune response. The balance between activation and inhibition is crucial for maintaining immune homeostasis and its dysregulation can lead to immune disorders. Thus, identifying new intracellular regulators of T cell signaling is particularly promising for designing new therapeutic strategies against cancer or autoimmune disorders ^4^. To find new components of the LAT-signalosome, we performed a proximity labeling assay of LAT and identified Microtubule Affinity-Regulating Kinase 2 (MARK2), a serine/threonine kinase of the AMPK family as part of the TCR signaling cascade ^5^. MARK2 has been primarily studied in the nervous system, where it regulates polarity, microtubule dynamics and cell migration ^6–9^ and its mutations are associated with neurological conditions such as autism spectrum disorders and other cognitive impairments ^10–12^. Outside the nervous system, MARK2 has been implicated in cancer progression and is known to be involved in several ubiquitous signaling pathways, such as WNT, PI3K-AKT-mTOR, and NF-κB ^13,14^. The role of MARK2 in the immune system remains poorly understood. Indeed, MARK2 is ubiquitously expressed and its germline deletion in mice results in a complex phenotype characterized by a dysregulation of T and B lymphocyte homeostasis, metabolic alterations, and signs of autoimmunity ^15^. These observations suggest a role for MARK2 in immune regulation, but its broad range of expression makes it difficult to determine its function in specific immune cell subsets.

To unravel the specific role of MARK2 in T cell biology we generated mice with a T-cell deletion of MARK2. We report here that MARK2 is required to maintain CD8 T cell homeostasis and acts as a negative regulator of sustained T cell activation in both murine and human cells. Moreover, we show that MARK2 controls the CD28-dependent co-stimulatory pathway and modulates the activation of the PI3K-AKT-mTORC1 signaling pathway as well as the glycolytic metabolism in T cells. Our data highlights a previously unappreciated role for this kinase in controlling co-activation and metabolic pathways in T cells, preventing excessive immune responses that could lead to autoimmunity.

## Results

### Generation of T-cell specific MARK2-deficient mice and thymocyte characterization

To study the role of MARK2 in T cell biology *in vivo*, we generated a conditional knockout mouse line in which exons 2 and 3 of *Mark2* were flanked by LoxP sequences (MARK2^flox/flox^ mice). These mice were crossed onto CD4-Cre transgenic mice to achieve a T-cell-specific deletion of MARK2 (Extended data Fig.1A). Immunoblotting analysis confirmed that MARK2 protein was almost undetectable in peripheral CD4 and CD8 T cells isolated from conditional KO (MARK2^cKO^) mice whereas its expression in B cells remained similar to wild type (WT) (Extended data Fig.1B). Of note, immunoblot quantification revealed that MARK2 protein level was approximately 1.3-fold higher in CD8⁺ T cells than in CD4⁺ T cells (Extended data Fig.1C).

**Figure 1.**
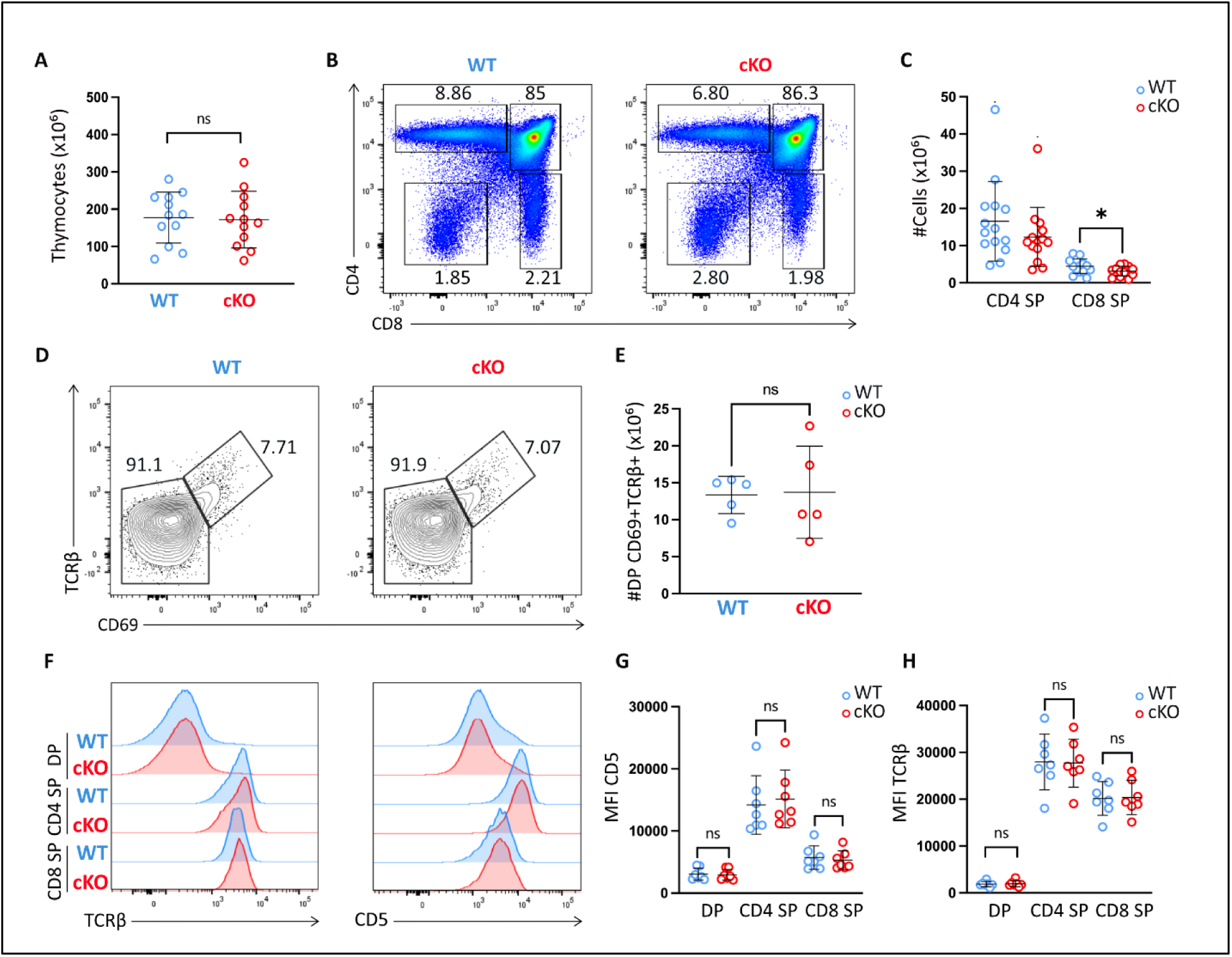
Thymic characterization of MARK2^cKO^ mice. Flow cytometry analysis of thymi harvested from wild-type (WT, in blue) or MARK2^cKO^ mice (in red)**. (A)** Total thymic cellularity**. (B)** Representative plots of CD4 and CD8 expression on live thymocytes isolated from WT or MARK2^cKO^ mice. **(C)** Absolute numbers of CD4^+^ and CD8^+^ single-positive (SP) thymocytes. **(D)** Representative plots of CD69 and TCRβ expression on double-positive CD4^+^ CD8^+^ (DP) cells from WT or MARK2^cKO^ mice. **(E)** Absolute numbers of positively selected (CD69^+^ TCRβ^+^) DP thymocytes. **(F)** Representative histograms showing CD5 and TCRβ expression on DP, CD4 SP and CD8 SP thymocytes from WT or MARK2^cKO^ mice, with quantifications shown in (**G)** and (**H)**. Each dot represents an individual mouse (n=14 mice for each genotype in **A-C**, n=5 in **E** and n=7 in **G-H**). All data were obtained from independent experiments. In graphs, bars represent mean and SD. Normality was assessed using the Shapiro-Wilk test. Statistical significance was determined by unpaired t-test for normally distributed data or, by Mann-Whitney test otherwise. In **C**, *P=0.0453.

Flow cytometry analysis of the thymus showed that MARK2 deficiency did not affect overall thymocyte numbers and subset differentiation (Fig.1A and B). We only observed a slight decrease in the absolute number of CD4^+^ and CD8^+^ single-positive (SP) thymocytes in the absence of MARK2 (Fig.1C). Expression of key developmental markers such as TCRβ, CD5 and CD69 was similar in MARK2^cKO^ and WT mice. The percentage of DP TCRβ^+^CD69^+^ cells, which represent positively selected T cells, was similar in KO and WT mice, suggesting that positive selection occurred normally in absence of MARK2 (Fig.1D to H).

Taken together, these data indicate that MARK2 is dispensable for the development and the positive selection of thymocytes *in vivo*.

### MARK2 regulates the number of central memory CD8+ T cells in a T-cell intrinsic manner

We next looked at T cell development in peripheral lymphoid organs to evaluate the role of MARK2 in mature T cell homeostasis. Spleens of 8–14 weeks old MARK2^cKO^ mice have the same number of total CD4 and CD8 T cells than WT littermates (Fig. 2A to C). However, a slight reduction in the number of CD8⁺ T cells was observed in the lymph nodes of MARK2^cKO^ mice (Extended Data Fig. 2A to C). To better understand whether MARK2 influences the development of T cell subsets, we analyzed by flow cytometry the spleen and LN of MARK2^cKO^ and WT mice stained for CD44 and CD62L. MARK2 deficiency did not strongly affect the number and the proportion of CD4^+^ T cell subsets in both spleen and LN (Fig.2D to F and Extended data Fig.2C). In contrast, we found a significant increase in both proportion and absolute number of CD44^hi^ CD62L^hi^ central memory (cM) CD8^+^ T cells in the spleen of MARK2^cKO^ mice, while CD44^low^CD62L^hi^ naïve CD8^+^ T cell proportion and number were concomitantly decreased compared to WT (Fig. 2G-H). A similar increase in central memory CD8⁺ T cells was observed in the lymph nodes (Extended data Fig.2D-E). Of note, expression levels of TCRβ and CD5, which correlate with the TCR avidity for self-peptide-MHC complexes, were similar in MARK2^cKO^ and WT mice (Extended data Fig. 2F to H). We noticed an increased expression levels of CD44 and CD62L in the naïve CD4⁺ and CD8⁺ T cell compartments of MARK2^cKO^ mice compared to WT (Fig.2I to M). A similar increase in CD44 in CD4⁺ T cells was previously reported in MARK2-deficient mice (MARK2^KO^), in which MARK2 was ubiquitously depleted. These results indicate that MARK2 deficiency alters the expression of key surface markers of T cell memory and trafficking. To further investigate whether the altered phenotype observed in MARK2^cKO^ mice were T cell intrinsic, we generated mixed bone marrow (BM) chimeric mice by injecting equal number of WT and MARK2^cKO^ BM cells into irradiated Rag2^KO^ mice (Extended data Fig.2I). Eight weeks after BM transfer, MARK2^cKO^ BM-derived CD8^+^ T cells showed a significant increase in the percentage of cM cells and elevated expression of CD44 and CD62L compared to WT BM-derived cells (Extended Data Fig. 2J to N), thus recapitulating the phenotype observed in MARK2^cKO^ mice. Taken together, these results indicate that MARK2 regulates the formation or maintenance of central memory CD8^+^ T cells in peripheral lymphoid organs in a T cell intrinsic manner but is dispensable for the homeostasis of the CD4+ subset.

**Figure 2.**
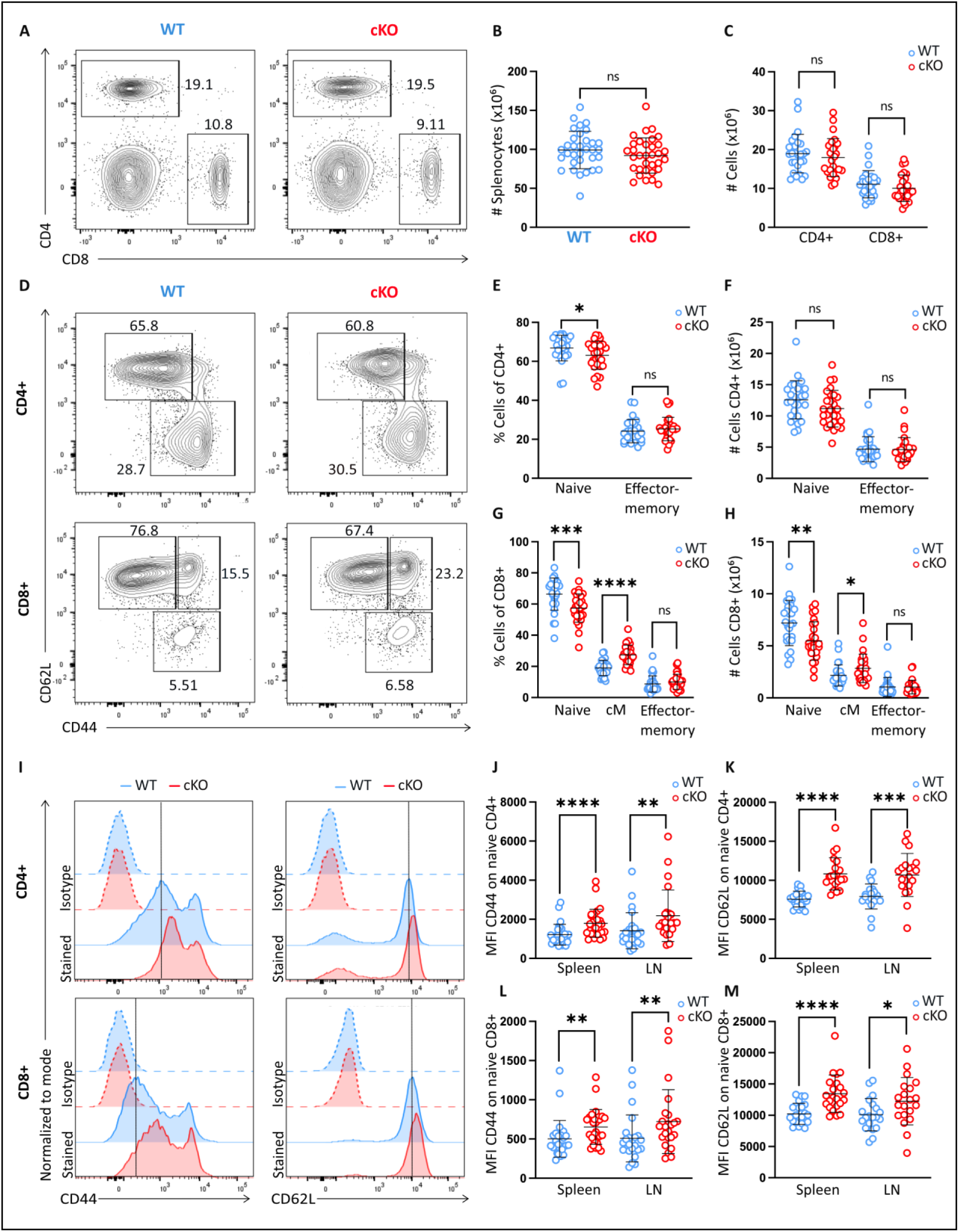
MARK2 regulates the formation of central memory (cM) CD8+ T cells in the spleen. **(A to G)** Flow cytometry analysis of spleens harvested from WT (in blue) or MARK2^cKO^ mice (in red). **(A)** Representative plots of CD4 and CD8 expression on live splenocytes isolated from WT or MARK2^cKO^ mice. Numbers in quadrant indicate proportions of cells. **(B)** Total splenic cellularity. **(C)** Absolute numbers of CD4^+^ and CD8^+^ T cells. **(D)** Representative plots of CD62L and CD44 expression on gated CD4^+^ or CD8^+^ splenocytes from WT or MARK2^cKO^ mice. Numbers in quadrant indicate proportion of naïve (CD44^lo^CD62L^hi^), effector memory (CD44^hi^CD62L^neg^) and central memory (CD44^hi^ CD62L^hi^) subsets. **(E, F)** Frequency and absolute numbers of each subset among CD4^+^ and **(G, H)** CD8+ T cells. **(I)** Representative histograms of CD44 and CD62L expression on gated CD4+ and CD8^+^ T cells. Dashed lines indicate isotype control levels. **(J, K)** Quantification of CD44 and CD62L expression on gated naïve CD4^+^ and **(L, M)** CD8^+^ T cells. Each dot represents an individual mouse (n=33 mice in **B**, n=27-28 in **C-H**, n= 20-28 in **J-M**). All data were obtained from independent experiments. In graphs, bars represent mean and SD. Statistical significance was determined by unpaired t-test for normally distributed data or, by Mann-Whitney test otherwise. In **E**, *P= 0,0371. In **G,** ***P = 0.0004, ****P < 0.0001. In **H**, **P=0.0027 and*P=0.0326. In **J,** **** P < 0.0001 and **P=0.0027. In **K**, ****P < 0.0001 and ***P=0.0004. In **L**, **P=0.0023 for spleens and **P=0.0081 for LN. In **M**, ****P < 0.0001 and *P=0.0421.

### MARK2 deletion in T cells leads to mild systemic autoimmunity upon aging

Hurov *et al.* described that MARK2^KO^ mice develop a late-onset autoimmune disease characterized by splenomegaly and/or lymphadenopathy in around 30% of aged mice, and immune infiltrates in non-lymphoid organs^15^. However, since MARK2 was ubiquitously depleted in that mouse model, it remains unclear which cells would be responsible for this immune dysregulation. To determine whether the specific deletion of MARK2 in T cells can trigger autoimmunity, we performed serological and histological analyses in MARK2^cKO^ and WT mice. As shown in Fig.3 A and B, we looked for the presence of anti-nuclear antibodies (ANA) in the serum of MARK2^cKO^ and WT mice aged 19 weeks to nearly 2 years. Compared with WT littermates, sera from aged MARK2^cKO^ displayed a characteristic nuclear staining pattern of HEP-2 cells and significantly elevated fluorescence intensity (referred to ANA score), indicative of increased anti-nuclear antibody levels. Furthermore, histological examination revealed marked lympho-plasmatic infiltrates in the kidneys, lungs, liver, and pancreas of aged MARK2^cKO^ mice, whereas these infiltrates were absent to mild in WT littermates. The histological score obtained from the semi-quantitative evaluation of tissue lesions was also found significantly higher in aged MARK2^cKO^ mice compared with WT littermates (Fig. 3C-D). Regulatory T cells (Tregs) play an important role in maintaining immune tolerance ^16^. We analyzed by flow cytometry Treg cells from MARK2^cKO^ and WT mice. We found no significant differences in the absolute number of CD25^+^Foxp3^+^ Treg in young (Fig. 3E-F) and old mice of both genotypes (Fig. 3G-H). Moreover, expression of key markers associated with Treg stability and suppressive functions such as CD25, Foxp3, GITR and PD-1, were similar between MARK2^cKO^ and WT littermates mice (Fig.3I-J). In contrast to what we previously described for conventional CD4^+^ cells, CD44 expression was not increased in Treg cells found in MARK2^cKO^ mice (Fig. 3K). Our data show that MARK2 acts to maintain peripheral immune tolerance, as its deletion in T cells leads to the spontaneous development of autoimmunity upon aging, independently of changes in Tregs phenotype and number.

**Figure 3.**
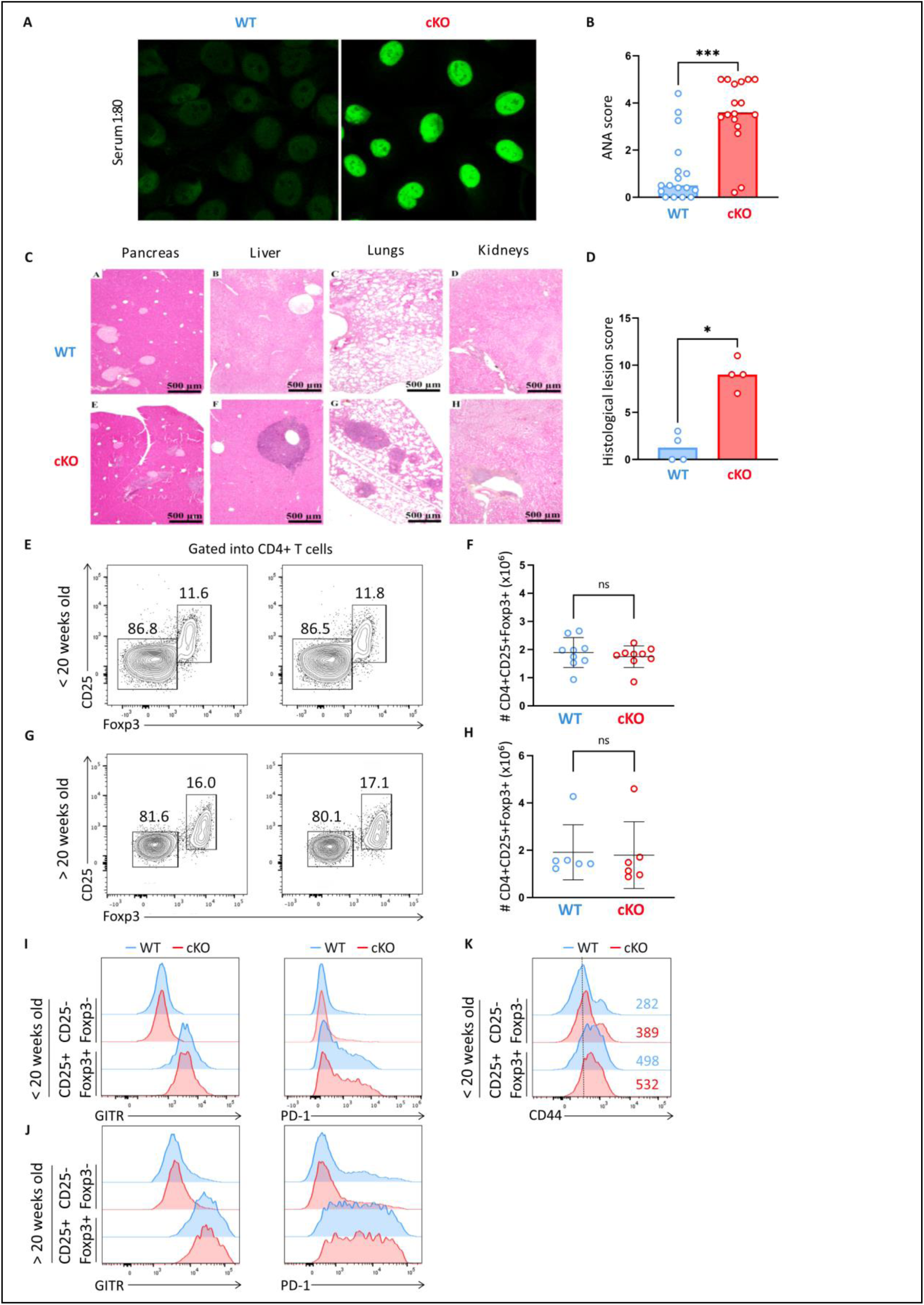
MARK2^cKO^ mice develop systemic autoimmunity upon aging. **(A)** HEP-2 cells were incubated with sera from aged WT or MARK2^cKO^ mice (19 weeks to 2 years old) and revealed with anti-mouse IgG-A488. The image shown is representative of the staining obtained with sera samples diluted at 1/80. **(B)** ANA score was quantified based on the fluorescence intensity obtained by microscopy. **(C)** Representative photographs of histological tissue sections of pancreas, liver, lungs and kidneys of WT and MARK2^cKO^ mouse littermate showing lympho-plasmatic infiltrates in organs, HES stain. **(D)** Quantification of histological lesion score. **(E to K)** Flow cytometry analysis of regulatory T cell (Tregs) found in the spleens of young and old (age >20 weeks) WT or MARK2^cKO^ mice. **(E, F)** Representative plots of CD25 and Foxp3 expression on gated CD4^+^ T cells and absolute numbers of CD4^+^CD25^+^Foxp3^+^ cells in young and **(G, H)** old mice. **(I, J)** Representative histograms showing GITR, PD-1 and **(K)** CD44 expression on gated conventional CD4^+^ T cells (CD25^-^Foxp3^-^) or Tregs (CD25^+^Foxp3^+^) from WT or MARK2^cKO^ mice. Geometric means are indicated. Each dot represents an individual mouse (n=17 mice in **B**, n=4 in **D**, n=9 in **F**, n=6 in **H**). Data were obtained from at least three independent experiments. In graphs, bars represent mean and SD. Statistical significance was determined by unpaired t-test for normally distributed data or, by Mann-Whitney test otherwise. In **B**, ***P=0.0002. In **D**, *P=0.0286.

### MARK2 is dispensable for early TCR signaling of antigen-stimulated CD8^+^ T cells

As CD8 T cell homeostasis was altered in MARK2^cKO^ mice, we then asked whether MARK2 deletion would affect TCR signaling in CD8^+^ T cells. We crossed MARK2^cKO^ mice with the TCR OT-I transgenic mice which express a TCR specific for an ovalbumin derived peptide presented by the MHC class I molecule H-2Kb (MARK2^cKO^ OT-I) ^17^. Splenocytes from MARK2^cKO^ OT-I mice exhibited the same elevated expression of CD44 and CD62L on naïve CD8^+^ T cells and increased proportion of central memory than we described for the MARK2^cKO^ mice on a polyclonal background (Extended data Fig.4A to E). Expression levels of OT-I TCR Vα2 and Vβ5 chains and of CD28 on CD8^+^ T cells, were comparable between MARK2^cKO^ and WT OT-I mice indicating similar TCR and costimulatory receptor availability (Extended data Fig.4F-G). We stimulated naïve CD8 T cells isolated from MARK2^cKO^ or WT OT-I mice in vitro with strong SIINFEKL (N4) or weak SIIGFEKL (G4) agonist peptides. In this assay, TCR stimulation can be evaluated independently of costimulatory signals provided by an antigen presenting cell (APC), allowing a direct readout of TCR signaling capacity. MARK2 deficiency had no effect on early TCR-induced activation responses such as CD69 and CD25 upregulation after overnight stimulation with different concentrations of both N4 and G4 peptides (Extended data Fig.4H to K). Furthermore, stimulation with N4/H-2Kb tetramers induced similar phosphorylation levels of ZAP70, PLCγ1 and S6 in MARK2^KO^ and WT OT-I T cells (Extended data Fig. 4L-M). Thus, MARK2 deficiency does not impact TCR signaling strength and early activation responses to peptide stimulation.

**Figure 4.**
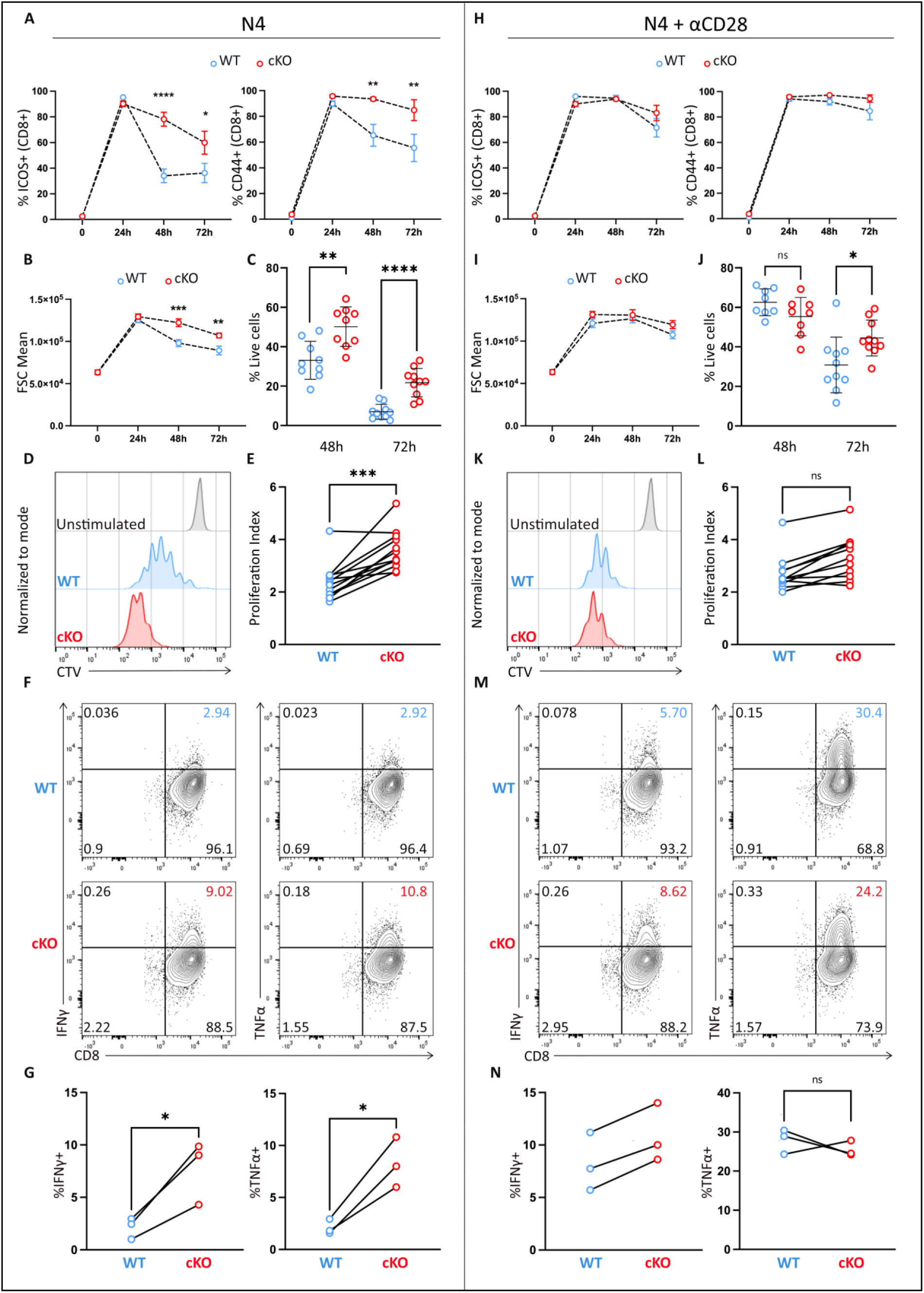
MARK2 deficiency in CD8 T cells leads to CD28 co-stimulation independence. Naïve CD8^+^ T cells were isolated from WT or MARK2^cKO^ OT-I mice and stimulated *in vitro* over time with 1nM of SIINFEKL (N4) peptide in the absence **(A to G)** or presence **(H to N)** of soluble anti-CD28 (1μg/ml). **(A, H)** Upregulation of CD44 and ICOS expression, **(B, I)** cell size and **(C, J)** viability, were quantified by flow cytometry at different time points following activation (as indicated on the x axis). **(D, K)** Representative histograms of CTV dilution in WT or MARK2^KO^ CD8^+^ T cells after 72h of activation compared to unstimulated cells (in gray). **(E, L)** Flow cytometry analysis of the proliferation index. **(F, M)** Representative plots showing IFNγ and TNFα expression in WT or MARK2^KO^ CD8^+^ T cells activated for 24h in presence of BrefeldinA for the last 4H. **(G, N)** Percentages of IFNγ and TNFα producing cells were quantified for each condition. In **A, B, H and I**, graphs show mean and SEM. Data were generated from n=5 independent experiments for **A and H,** and n=7 independent experiments for **B and I**. Statistical significance was determined using two-way ANOVA. In **A**, **** P<0.0001 and *P=0.0114 for ICOS, **P=0.0074 and **P=0.0052 for CD44. In **B**, ***P=0.0001 at 48h and **P=0.0055 at 72h. In **C, E, G, J, L and N**, each dot represents an individual mouse and data were obtained from at least n=3 independent experiments. Statistical significance was determined by unpaired t-test for normally distributed data or, by Mann-Whitney test otherwise. In **C,** **P=0.0020 and ****P < 0.0001. In **E**, ***P=0.0001. In **G**, *P=0.0330 for IFNγ and *P=0.0178 for TNFα. In **J**, *P=0.0192.

### MARK2 regulates sustained TCR-induced activation of CD8^+^ T cells

We next evaluated late activation events after 24h, 48H, or 72H of *in vitro* TCR stimulation. While ICOS and CD44 expression levels were comparable at 24 hours in both genotypes, their expression in WT cells declined at later timepoints, whereas MARK2-deficient T cells maintained high ICOS and CD44 levels through 72 hours (Fig. 4A). MARK2^KO^ OT-I cells also exhibited increases in cell size (FSC) and survival over time (Fig. 4B-C). Next, we tested the capacity of MARK2-deficient OT-I T cells to sustain effector functions in response to N4 peptide stimulation. As shown in Fig. 4D and E, the proliferation of MARK2^KO^ OT-I cells was enhanced compared to WT after 3 days of stimulation, as shown by a significant increase in the proliferation index. Furthermore, after 24H of N4 stimulation, the percentage of CD8^+^ cells producing TNFα or IFNγ were significantly increased in MARK2^KO^ OT-I cells compared to WT cells (Fig. 4F-G). Importantly, both genotypes responded similarly to PMA/ionomycin stimulation, which bypasses proximal TCR signaling, indicating that MARK2 does not affect distal signaling events (Extended Data Fig. 4 bis A to C). To determine whether the enhanced proliferation and cytokine production in MARK2^KO^ T cells was driven by a cytokine mediated feedback loop, we incubated WT T cells with supernatant derived from MARK2^KO^ T cell culture.

This had no significant impact on the activation, survival, or proliferation of WT OT-I T cells (Extended data Fig.4 bis D to F), indicating that the sustained activation phenotype induced by MARK2 deficiency was a cell-intrinsic mechanism and was not driven by increased cytokine secretion. Altogether these results show that MARK2 is not involved in early CD8 T cell responses to antigenic stimulation, but limits sustained activation and effector functions induced by TCR triggering.

### MARK2 deficiency leads to CD28 co-stimulation independence

We have shown that MARK2 deficiency leads to sustained activation, enhanced survival and effector functions in response to antigen specific stimulation alone, processes that are normally dependent on CD28-mediated co-stimulation. To test whether this phenotype was due to a deregulation in the CD28 pathway, we stimulated naïve MARK2^KO^ and WT OT-I cells with N4 peptide in the presence of an agonistic anti-CD28 antibody. As expected, CD28 stimulation amplified and sustained the upregulation of activation markers, survival, cell size, proliferation, and cytokine production in WT OT-I T cells, reaching levels comparable to those observed in MARK2^KO^ OT-I cells. In contrast, CD28 co-stimulation had little to no additional effect on MARK2^KO^ OT-I cell activation (Fig.4H to N).

This CD28-independent phenotype was also observed in polyclonal naïve MARK2^KO^ CD8^+^ T cells stimulated with plate-bound anti-CD3 antibody as demonstrated by enhanced survival, cell size, activation and proliferation over time compared to CD8^+^ WT cells and, similar responses in both genotypes upon CD28 engagement (Extended data Fig.4 bis G to L).

To determine whether this phenotype was conserved in human T cells, we inactivated MARK2 by CRISPR-Cas9 in primary T cells isolated from human PBMCs (schematic representation of protocol in extended data Fig. 4 bis M and western-blot in Extended data Fig.4 bis N). Upon CD3 stimulation alone MARK2 inactivated T cells showed enhanced proliferation (Extended Fig.4 bis O-P) compared to control cells. However, in the presence of anti-CD3 and anti-CD28 antibodies, both MARK2 KO and control T cells display similar proliferation rates (Extended data Fig.4 bis O-P).

Altogether, our data show that MARK2-deficient T cells do not require CD28 triggering to sustain activation and functions thus suggesting that MARK2 is a key regulator of CD28-dependent co-stimulation.

### Sustained activation of MARK2^KO^ T cells is independent of CD28 co-receptor engagement with its ligands

Our results suggest that MARK2 is a regulator of the CD28 costimulatory pathway. It was recently shown that CD80 and CD86 can be induced upon TCR stimulation on T cells and can provide efficient co-stimulatory interactions with CD28 *in cis* ^18^. We thus wondered whether the sustained activation of MARK2-deficient T cells could be due to an increased expression of B7 molecules, which would provide CD28 engagement *in cis*. We looked at the upregulation of CD80 and CD86 by flow cytometry on naïve MARK2^KO^ and WT OT-I T cells primed with N4 peptide alone. MARK2-deficient T cells expressed more CD80 and CD86 molecules following TCR activation than their WT counterparts (Fig. 5A to D), while surface expression of CD28 remained similar (Fig.5E). Yet, increased proliferation of MARK2^KO^OT-I T cells was not due to this increased expression of CD80/86 since blocking the CD28/B7 interactions with CTLA4-Ig had little to no effect on proliferation of MARK2^KO^OT-I T cells while inhibiting WT T cell proliferation (Fig. 5F-G). These results indicate that sustained activation of MARK2^KO^ T cells occurs independently of CD28 engagement.

**Figure 5.**
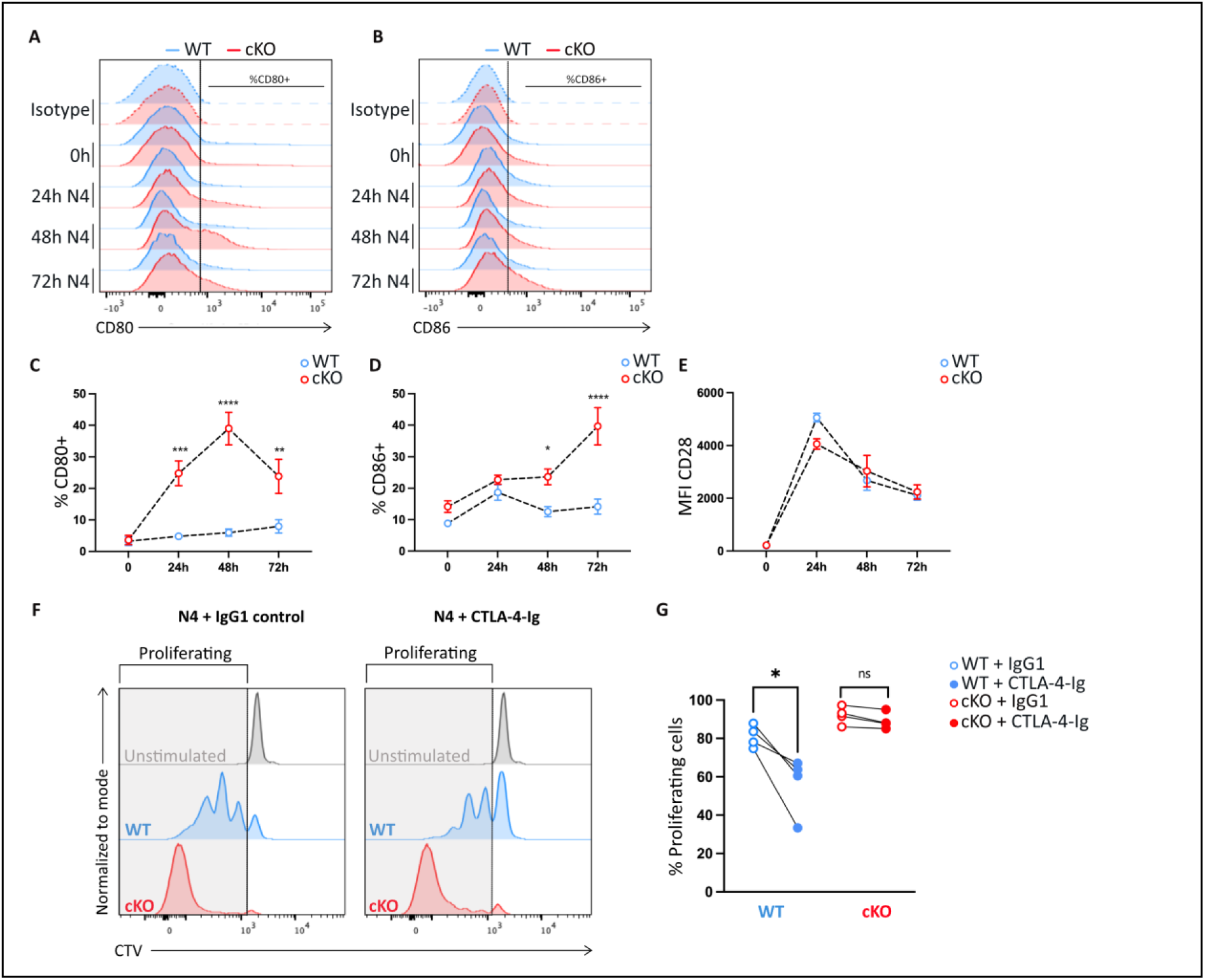
MARK2 ^KO^ T cell sustained activation is independent of CD28 co-receptor engagement with its ligands. Naïve CD8^+^ T cells isolated from WT or MARK2^cKO^ OT-I mice were stimulated *in vitro* with 1nM of N4 peptide for 24, 48 and 72 hours. **(A to B)** Representative histograms showing **(A)** CD80 and **(B)** CD86 expression at different time points following TCR stimulation (as indicated on the y axis). **(C to E)** Frequency of **(C)** CD80^+^**, (D)** CD86^+^ and **(E)** CD28^+^ cells among CD8^+^ T cells following activation. **(F)** Representative histograms of CTV dilution in WT or MARK2^KO^ CD8^+^ T cells after 72h of TCR stimulation in the presence of CTLA-4-Ig or IgG1 control (5μg/ml). Gray boxes represent proliferating cells and were quantified in **(G)** for each condition. In **C, D and E,** graphs show mean and SEM and were generated from at least n=4 independent experiments. Statistical significance was determined using two-way ANOVA. In **C**, ***P=0.0004, ****P < 0.0001 and **P=0.0046. In **D**, *P=0.0164 and ****P < 0.0001. In **G**, data were generated from n=4 independent experiments and statistical significance was determined using unpaired t-test. *P=0.0236.

### Single-cell RNA sequencing (sc-RNA seq) identifies distinct naïve clusters in MARK2^KO^ and WT T cells

To define the transcriptional impact of MARK2 deficiency in CD8 T cells, CD8α^+^, TCRβ^+^, CD62L^+^, CD44^low^ cells were isolated by FACS from MARK2^cKO^ or WT mice and subjected to single-cell RNA-sequencing (scRNA-seq). Cells were analyzed either unstimulated (T0, n=11667 cells) or following 5 or 24 hours of TCR activation (T5, n=10206 cells and T24, n=10043 cells) (Extended data Fig.6A). A Uniform Manifold Approximation Projection (UMAP) representation encompassing all samples revealed distinct cell populations that varied with the time of TCR stimulation (Fig.6A and Extended data Fig.6B). Unsupervised clustering of the datasets at resolution 0.2 (Extended data Fig.6C) identified five main clusters. At T0, two *naïve-like* clusters (called *naïve-like 1* and *naïve-like 2*) were identified both enriched in stemness-related genes such as *Il7r, Sell and Tcf7* (Fig.6B to D). Naïve-like 2 cluster expressed higher levels of these stemness-related genes compared to the naïve-like 1 cluster. Five hours after TCR activation, one cluster was over-represented and was enriched for early activation genes such as *Nr4a1* and *Egr1*. After 24h, two clusters appeared, one expressing proliferation (*Top2a*, *Mki67*) and activation (*Cd44*, *Pdcd1)* genes corresponding to the *proliferating cells*, and another that starts expressing *CD44* and *Top2a.* This latter cluster was called *early proliferating cells* since the cells started to down regulate stemness related genes and to upregulate activating and proliferating genes (Fig.6B to D). We next compared clusters proportion between genotypes and noticed some significant differences between MARK2^KO^ and WT T cells (Fig. 6E-F). While the proportion of naïve cells was almost similar in both genotypes (52% of total cells for the WT and 56% for the MARK2^KO^), their distribution within the naïve-like 1 and naïve-like 2 clusters was different. We found that at T0, naïve-like 2 cells were over-represented in MARK2-deficient cells (94% in MARK2^KO^ versus 6% in WT) while naïve-like 1 cells were strongly reduced (4% cells in MARK2^KO^ versus 93% in WT) (Fig. 6G). We also noticed that the naïve-like 1 cluster disappeared after TCR stimulation, whereas the naïve-like 2 cluster persisted even 24h post stimulation. Of note, the naive-like 2 cluster was still more represented in activated MARK2^KO^ T cells than in WT T cells (Fig. 6G). We also found an increased proportion of cells in the proliferating cluster in MARK2-deficient T cells (5% in the WT vs 8% in the KO), which was consistent with our previous observations (Fig. 6F). The increased expression of CD62L (encoded by *Sell*) observed by flow cytometry in MARK2-deficient naïve cells was also detected at the transcriptional level with *Sell* more expressed in the naïve-like 2 cluster than in the naïve-like 1 (see *Sell* in Fig 6C).

**Figure 6.**
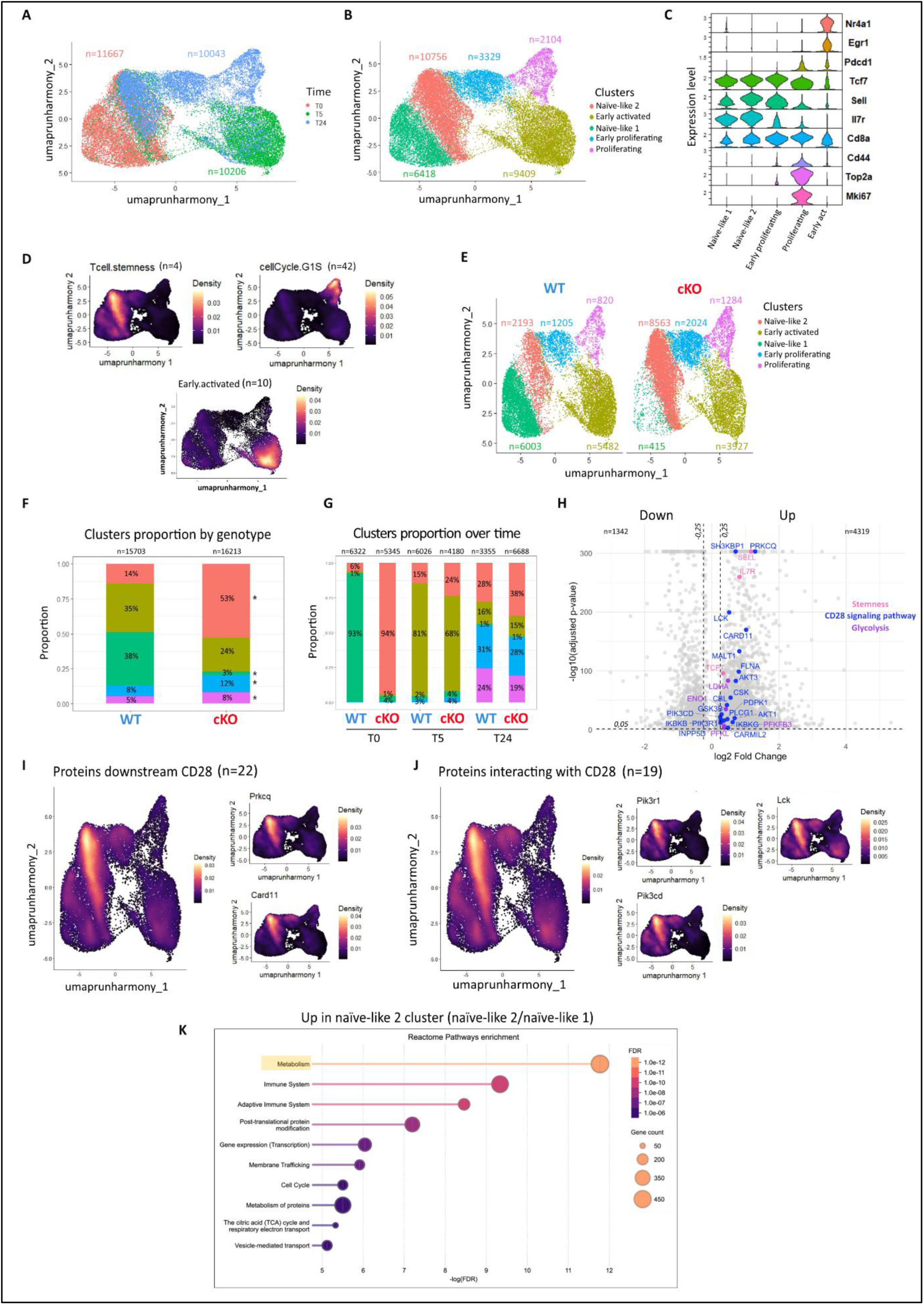
Single-cell RNA sequencing (sc-RNA seq) analysis revealed an enriched population of naïve-like cells expressing metabolism and CD28-related genes in MARK2 ^KO^ CD8^+^ T cells. Naïve CD8^+^ T cells from WT or MARK2^cKO^ mice were left unstimulated (T0) or stimulated *in vitro* with plate-bound anti-CD3 (5μg/ml) for 5 (T5) or 24 hours (T24) before RNA sequencing. **(A)** UMAP representations showing cell clustering by TCR activation time points and **(B)** the five main clusters identified. Cell counts in each cluster are indicated on the UMAP.**(C)** Violin plot showing expression of selected genes in each cluster and **(D)** UMAP representation showing specific gene signatures including *Tcf7, Sell* and *Il7r* in the “T cell stemness” signature, *Mki67* in the “Cell cycle G1-S” signature and *Nr4a1* and *Egr1* in the “early activated” signature. **(E to F)** UMAP representation and bar plots showing cluster distribution by genotype. Cell counts are indicated on the UMAP. **(G)** Bar plots depicting cluster proportion by genotype and across the different time points of TCR stimulation. Total cell counts in each condition are indicated on the top of the bars. **(H)** Volcano plot showing the genes upregulated in naïve-like 2 cluster compared to naïve-like 1 cluster. Genes associated with stemness (pink), CD28 signaling pathway (blue) and glycolysis (purple) are highlighted in the plot. Dashed lines indicated the parameters used for the analysis (adjusted p-value < 0,05 and log2 Fold Change set to 0,25). **(I to J)** UMAP visualizations of **(I)** “proteins downstream CD28” signature containing genes such as *Prkcq* and *Card11* and **(J)** “proteins interacting with CD28” signature including genes like *Pik3r1*, *Pik3cd* and *Lck*. **(K)** Top10 Reactome pathways obtained from the differential expression analysis of naïve-like 2 cluster vs naïve-like 1 cluster. The size of the circles represents the number of genes enriched in the pathway, and the color indicates the adjusted p-value. In **F**, statistical significance was determined by a permutation test. * was indicated if FDR<0,05 and abs(log2FD)>0,58.

### MARK2 controls transcriptomic pathways related to CD28 and glycolysis

We then further characterized the differences between the naive-like 1 and naive-like 2 clusters. A volcano plot representation comparing all the genes differentially regulated in the naive-like 2 versus naive-like 1 clusters identified 4319 genes up-regulated and 1342 genes down-modulated (cut off Log2FC= 0.25 and adjusted p-value < 0,05). Among the up-regulated genes, we found some genes of the stemness signature, genes belonging to the CD28 signaling pathways and genes belonging to the glycolysis pathway such as *Ldha*, *Eno1*, *Pfkfb3* and *Pfkl* (Fig. 6H). To deeply characterize the nature of the genes related to the CD28 pathway, we designed two gene signatures based on literature ^3,19^. The “Proteins downstream CD28” signature contained genes such as *Prkcq* and *Card11* and the “Proteins interacting with CD28” signature contained genes coding for proteins interacting with the CD28 cytoplasmic tail such as subunits of the PI3K, the p85 (*Pik3r1)* and the p110δ (*Pik3cd),* and others like *Lck*. Both gene signatures showed a striking enrichment in the naïve-like 2 cluster compared to the other clusters (Fig.6I-J). Together with the fact that the naive-like 2 cluster is overrepresented in MARK2^KO^ T cells, these results show that at steady state MARK2-deficient naïve CD8 T cells have a transcriptional profile enriched for genes involved in the CD28 signaling pathway. This can explain the sustained activation and increased proliferation of MARK2^KO^ T cells stimulated by the TCR alone. Finally, to find out which molecular pathways were related to the two identified naïve-like clusters, we performed a functional enrichment analysis using the Reactome database (Fig.6K). The pathways “metabolism” and “immune system” were the most enriched in the naïve-like 2 cluster compared to the naïve-like 1 cluster. This is consistent with one of the roles of CD28 costimulation which is to promote rapid up-regulation of glycolysis ^20^.

Altogether, these results indicate that MARK2 controls transcriptional programs involved in the CD28 co-stimulation and metabolic pathways.

### MARK2 regulates PI3K-AKT-mTOR activation pathway and glycolysis metabolism in T cells

To further demonstrate that the CD28 signaling pathway was deregulated in MARK2^KO^ naïve T cells, we analyzed mTORC1 activity, which plays a key role in the sustained activation induced by CD28 co-stimulation, by assessing phosphorylation of the S6 ribosomal protein (Ser^235/236^) after TCR stimulation^21^. Naïve OT-I cells were activated for 24h with the N4 peptide and S6 phosphorylation was detected by immunoblot. We observed a significant increase in the pS6 levels in MARK2^KO^ OT-I T cells compared with WT OT-I cells (Fig.7A-B). This increase was dependent on mTOR and PI3K activities since Rapamycin (an mTOR inhibitor) and Ly294002 (a PI3K inhibitor) strongly inhibited S6 phosphorylation (Fig. 7A and F). Of note, CD28 co-stimulation induced increased S6 phosphorylation in WT OT-I T cells, to levels comparable to those observed in MARK2-deficient T cells, but co-stimulation had no effect in MARK2^KO^ T cells (Fig. 7A-B and flow cytometry analysis Fig. 7C and F). We also detected a slight, but reproducible, increase in the phosphorylation of AKT at T308, downstream of the PI3K-PDK1 axis ^22^, in MARK2-deficient T cells activated by the TCR only. CD28 co-stimulation induced similar levels of AKT phosphorylation in both MARK2-deficient and WT T cells (Fig. 7D and G). We also tested the phosphorylation of the p65 subunit of NFκB at S529, which is not increased by CD28 costimulation, and found that it was similar in MARK2^KO^ and WT OT-I T cells in both conditions (Fig.7E and H). This suggests that MARK2 specifically controls the PI3K-AKT-mTORC1 axis downstream of CD28. The role of mTOR in coordinating naïve T cell differentiation into effector T cells has been attributed, in part, to its ability to promote glycolysis. To evaluate if MARK2 could regulate CD8 T cell metabolic pathways, we performed a glycolytic assay using the Seahorse technology. As shown in Fig.7I to K, we observed an increase in the basal glycolysis of MARK2^KO^ OT-I T cells compared to WT OT-I cells, 24h after TCR stimulation. As expected, CD28 engagement enhanced T cell basal glycolysis in WT OT-I T cells at the same level as MARK2^KO^ OT-I T cells. Consistent with our previous observations, CD28 triggering had only a little effect on the basal glycolysis of MARK2-deficient T cells. Since our sc-RNA seq analysis suggested that MARK2 could also regulate the metabolism of resting CD8^+^ T cells, we analyzed the mitochondrial dependance (Fig.7L) and the glycolytic capacity (Fig.7M) of resting naïve T cells by flow cytometry using the SCENITH method ^23^. We observed an increased glycolytic capacity of naïve CD8^+^ T cells at resting state in absence of MARK2. These results further support a regulatory role for MARK2 in CD8^+^ T cell metabolic programming.

**Figure 7.**
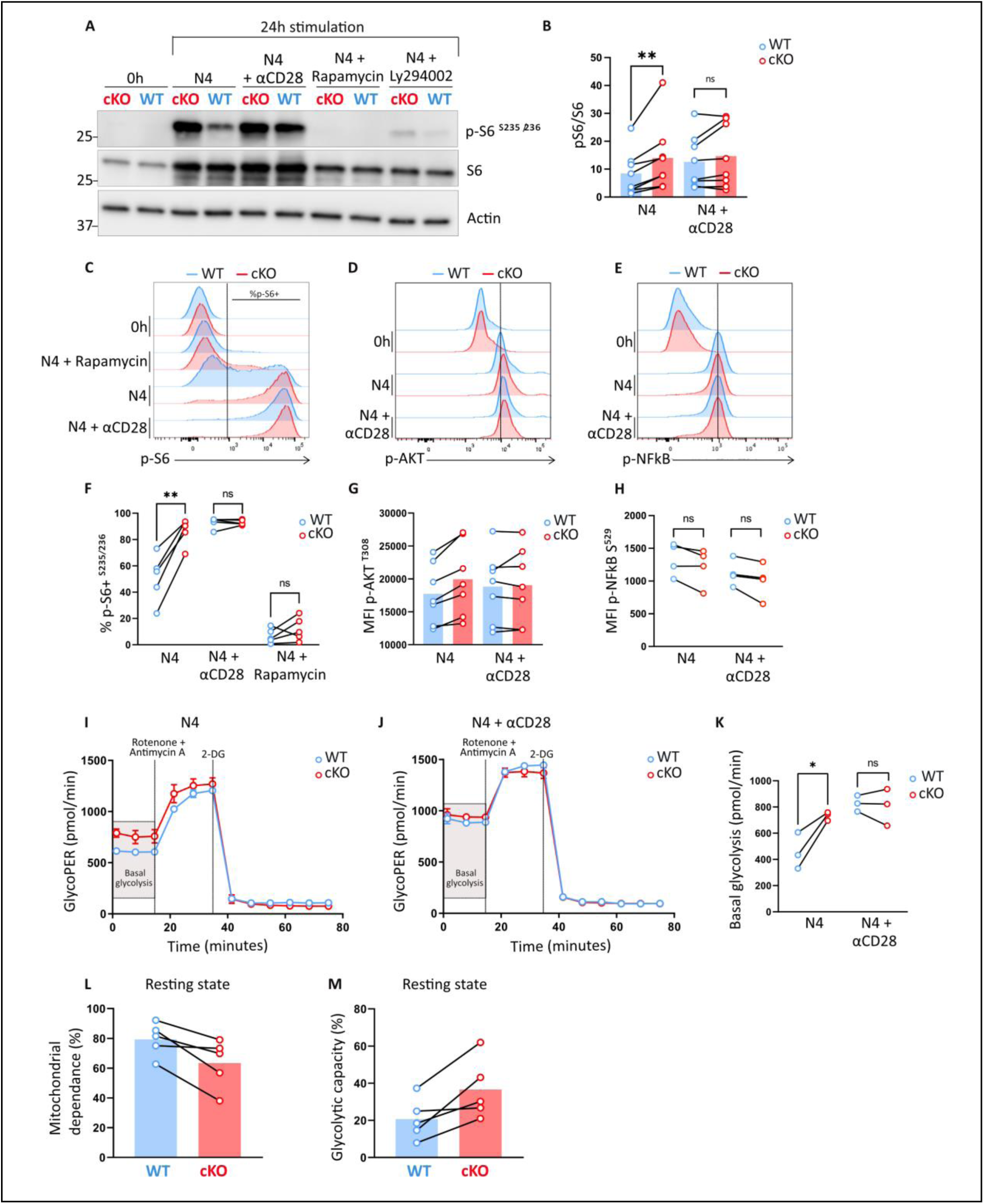
MARK2 regulates PI3K-AKT-mTOR activation pathway and glycolysis metabolism in T cells. **(A)** Representative immunoblot of total protein lysates from WT or MARK2 ^KO^ naïve CD8+ OT-I T cells stimulated for 24h with N4 peptide (1nM), or with N4 and soluble anti-CD28 (1μg/ml), rapamycin (20nM) or Ly294002 (10μM). Blot was probed with antibodies recognizing phospho-S6 (Ser^235/236^), S6 total and actin **(B)** Quantification of phospho-S6 band intensity relative to unstimulated WT cells band intensity and normalized to total S6 protein levels. **(C to H)** Phospho-flow cytometry showing **(C)** phospho-S6 (Ser^235/236^), **(D)** phospho-AKT (T^308^) and **(E)** phospho-NFκB p65 (S^529^) levels in WT or MARK2 ^KO^ naïve CD8^+^ OT-I T cells stimulated for 24h with N4 peptide alone or with anti-CD28 or rapamycin (when indicated). **(F)** Frequency of phospho-S6+ cells (compared to rapamycin treated condition) and MFI of **(G)** phospho-AKT and **(H)** phospho-NFκB were quantified for each condition. **(I to K)** Glycolysis assay using Seahorse technology performed in WT or MARK2 ^KO^ naïve OT-I T cells stimulated for 24h with N4 peptide **(I)** alone or **(J)** in presence of anti-CD28. Gray boxes indicate basal glycolysis levels and were quantified in **(K)** for each condition. **(L to M)** Flow cytometry analysis using SCENITH technology of **(L)** the mitochondrial dependance and **(M)** the glycolytic capacity of unstimulated WT or MARK2 ^KO^ naïve CD8^+^ OT-I T cells. Each dot represents an individual mouse (n=8 mice in **B**, n=4-7 mice in **F to H**, n=3 in **K**, n=5 in **L-M**). Data were generated at least from n=3 independent experiments. In **I and J**, graphs show mean and SEM from three technical replicates. Normality was determined by the Shapiro-Wilk test. In **B**, statistical significance was calculated using a paired t-test **P=0.025. In **F and K**, statistical significance was determined using an unpaired t-test. In **F**, **P=0.0054. In **K**, *P=0.0287.

### MARK2 regulates CRTC2 phosphorylation and CREB-dependent signaling in T cells

Although MARK2 has been implicated in many signaling pathways, its downstream targets in T cells remain uncharacterized. In pancreatic cells, MARK2 has been identified as a kinase phosphorylating the CREB-regulated-transcriptional-coactivator CRTC2 specifically at Ser275^24^. In this context, phosphorylation of CRTC2 promotes its interaction with 14-3-3 proteins leading to its cytoplasmic localization and preventing its interaction with CREB in the nucleus and the subsequent expression of CREB-related genes. Moreover, in cancer cells, CREB has been shown to regulate the expression of genes related to the PI3K-AKT axis, such as the p85 subunit of PI3K ^25^. To investigate whether this regulatory axis also occurred in T cells, we analyzed CRTC2 phosphorylation at Ser275 in WT and MARK2^KO^ CD8⁺ OT-I T cells stimulated for 24 hours with the N4 peptide, with or without CD28 co-stimulation. TCR stimulation induced CRTC2 phosphorylation at Ser275 in WT cells, which was reduced in MARK2^KO^ T cells (Fig.8A-B). Upon CD28 co-stimulation, CRTC2 phosphorylation was reduced in WT cells, suggesting that CD28 signaling inhibited MARK2 kinase activity on CRTC2. Treatment with rapamycin increased Ser275 phosphorylation in both WT and MARK2-deficient T cells, suggesting that the mTOR pathway is involved in the regulation of CRTC2 phosphorylation (Fig.8A-B). These data demonstrate that MARK2 promotes the specific phosphorylation of CRTC2 on Ser275 upon TCR stimulation in CD8 T cells, which is negatively regulated by CD28 costimulatory signals. To further explore the link between MARK2, mTOR and the CRTC2-CREB pathway, we assessed the impact of CREB inhibition on S6 phosphorylation in MARK2^KO^ or WT OT-I CD8^+^ T cells by flow cytometry (Fig.8C to E). The pharmacological inhibition of CREB strongly reduced pS6 levels in both MARK2-deficient and WT T cells, showing a regulatory feedback loop between CREB and the mTORC1-S6 axis (see N4+CREBi in Fig.8C). Finally, to evaluate CREB activity in WT and MARK2^KO^ CD8 T cells, we performed a CRE-luciferase reporter assay. Our data (Fig.8F) show that CREB-transcriptional activity was increased in absence of MARK2 compared to WT cells both upon N4 and N4+CD28 activation.

**Figure 8.**
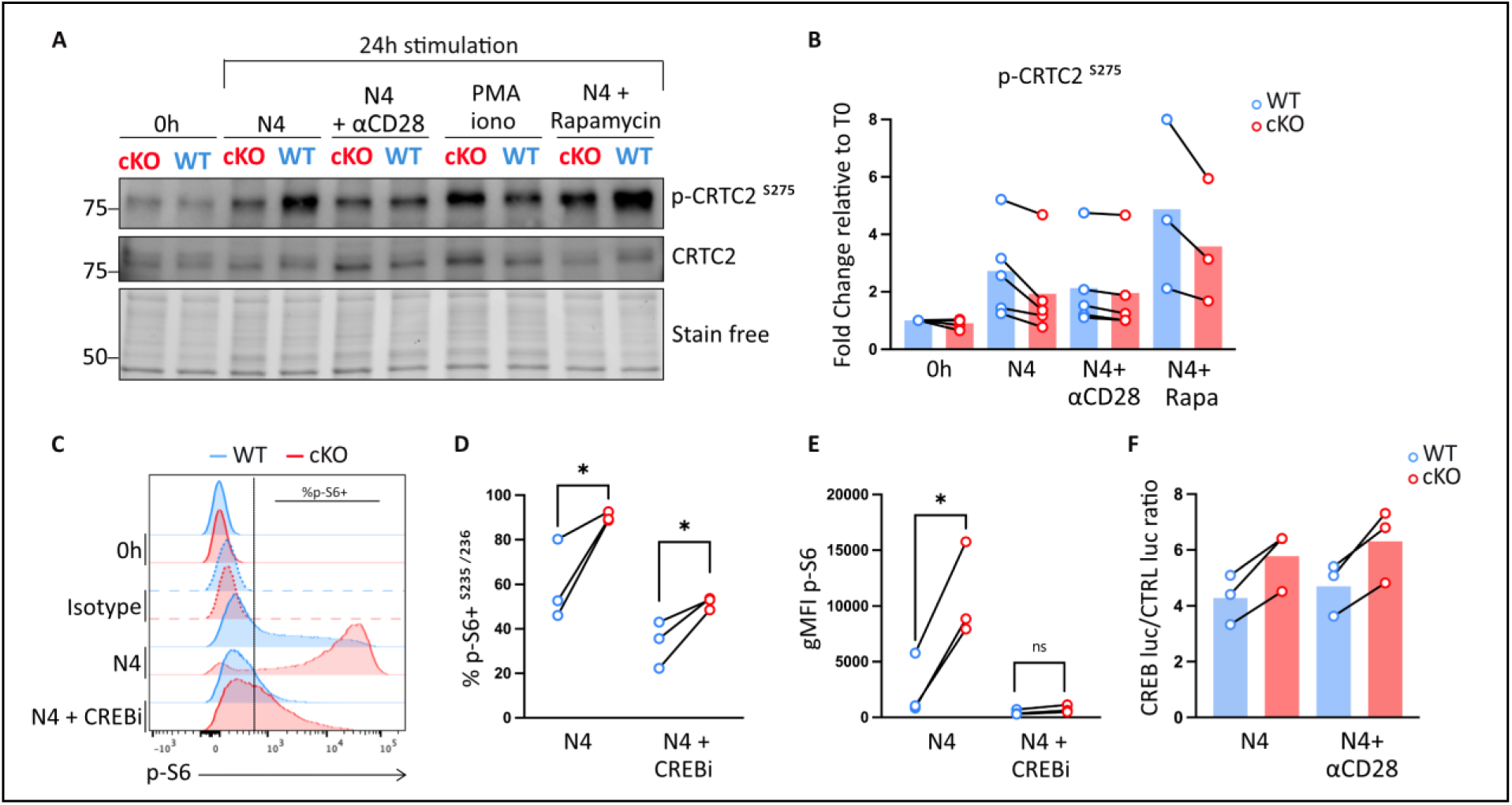
MARK2 controls CRTC2 phosphorylation and CREB-dependent signaling in CD8 T cells. **(A)** Representative immunoblot of total protein lysates from WT or MARK2 ^KO^ naïve CD8+ OT-I T cells stimulated for 24h with N4 peptide (1nM), either alone or in presence of soluble anti-CD28 (1μg/ml), PMA-ionomycin or rapamycin (20nM). Immunodetection was performed using antibodies specific for phospho-CRTC2 (Ser^275^) and total CRTC2. Total protein levels (stain free) were used as a loading control. **(B)** Quantification of phospho-CRTC2 band intensity relative to unstimulated WT cells band intensity and normalized to stain free. **(C to E)** Flow cytometry analysis of **(C)** phospho-S6 (Ser^235/236^) expression in WT or MARK2^KO^ naïve CD8^+^ OT-I T cells stimulated for 24h with N4 peptide alone or in combination with the CREB inhibitor 666-15 (5μM). **(D)** Frequency of phospho-S6+ cells, as indicated in **C**, and **(E)** geometric MFI of phospho-S6 were determined for each condition. **(F)** CREB activity measured by a CRE-inducible luciferase assay in CD8+ OT-I cells stimulated post-transfection for 24hours with N4 or N4+CD28. Quantification showing the ratio of induced CREB-luciferase value/ constitutive luciferase control expression. Each dot represents an individual mouse (n=3-5 mice in **B**, n= 3 in **D-F**). Data were generated from at least n=3 independent experiments. In **D-E**, statistical significance was obtained using an unpaired t-test. In **D**, *P=0.0443 for N4 and *P=0.0449 for N4+CREBi. In **E**, *P=0.0481.

**Figure 9.**
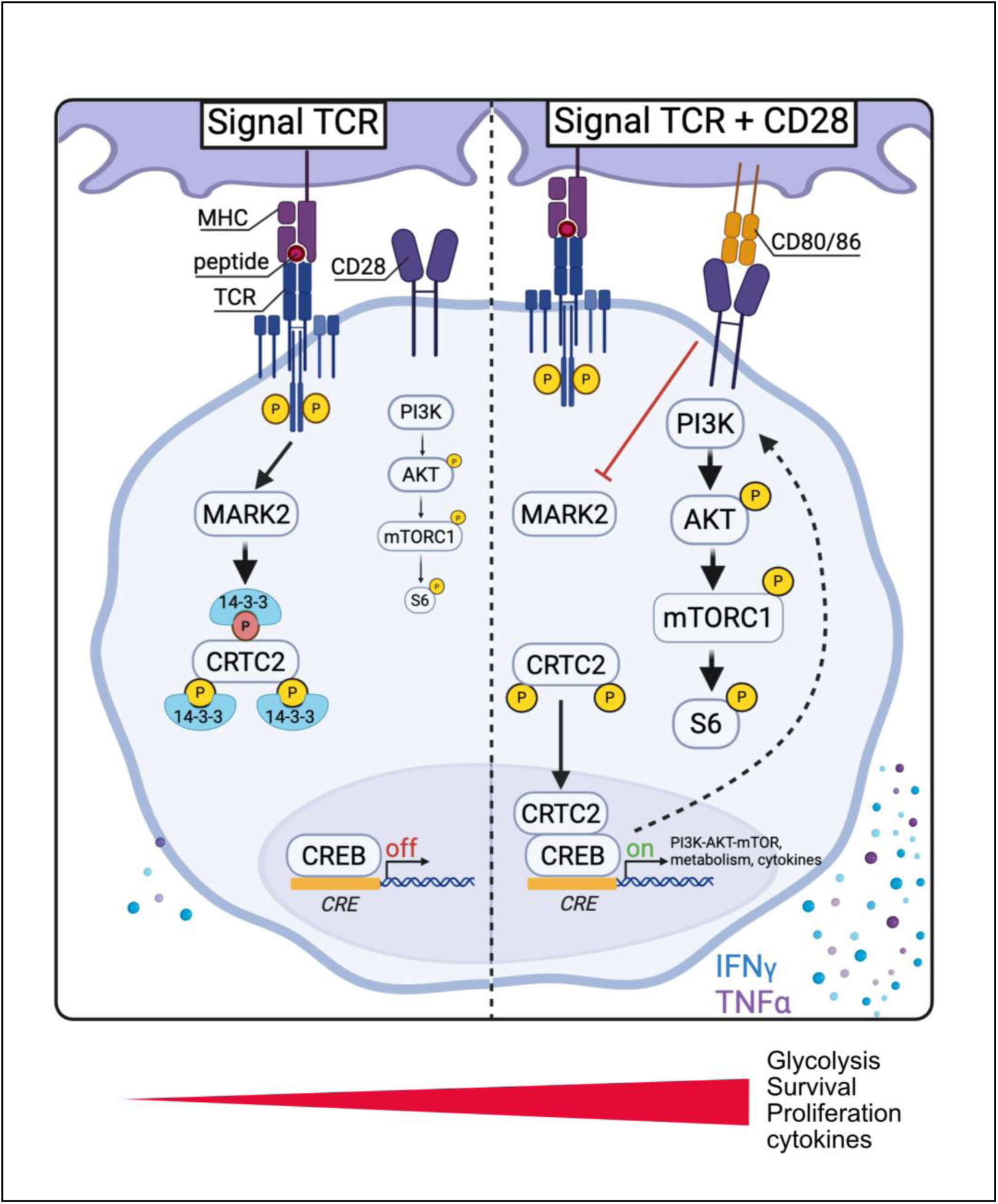
Graphical abstract. Naïve T cells express both the TCR and CD28 at their surface however, under resting conditions, antigen presenting cells do not express CD80/CD86 which are the ligands for CD28. TCR engagement in absence of CD28 co-stimulation leads to the activation of the kinase MARK2 which then phosphorylates CRTC2 at Ser275. This phosphorylation promotes CRTC2 sequestration in the cytoplasm through binding to 14-3-3 proteins. As a result, CRTC2 is not able to serve as a CREB coactivator resulting in an inhibition of CREB transcriptional activity. This inhibitory signal contributes to the induction of cell death, thereby preventing inappropriate T cell activation in the absence of a “danger signal”. When a pathogen enters the body, antigen presenting cells are activated and up-regulate CD80 and CD86. Upon co-engagement of the TCR and CD28, MARK2-mediated phosphorylation of CRTC2 on S275 is relieved, allowing CRTC2 to translocate to the nucleus and to promote the expression of CREB target genes. Some of the CREB-regulated genes control the PI3K-AKT-mTOR pathway and glucose metabolism, supporting full T cell activation and effector functions. In absence of MARK2, we observe a diminution of CRTC2 phosphorylation and an increased CREB activity upon stimulation by the TCR alone. This induces an overactivation of the PI3K-AKT-mTOR pathway which can explain why MARK2-deficient T cells have sustained proliferation, survival, glycolysis, and cytokine secretion. This mechanism ensures that TCR signaling alone is not sufficient to induce full T cell activation without a co-stimulatory “danger signal” and thus protecting from autoimmunity. Our findings show that MARK2 is a central checkpoint that prevents ligand-independent CD28 signaling and limits aberrant T cell activation.

Collectively, these results identify MARK2 as a key regulator of the CRTC2-CREB axis in TCR-stimulated CD8 T cells. By specifically phosphorylating CRTC2 at S275, MARK2 prevents CREB transcriptional activity and may control the expression of genes related to the PI3K-AKT-mTORC1 axis and to glucose metabolism.

## Discussion

Although the MARK2 kinase was cloned almost 30 years ago, ^26^ and full knockout (KO) mice have been available for many years, its role in the immune system has remained poorly studied ^15,27,28^. A 2001 study reported that MARK2^KO^ mice had normal B and T cell development but displayed signs of immune dysregulation, including increased CD44 expression on CD4⁺ T cells, elevated cytokine production upon TCR engagement, and enhanced antibody responses to T cell–dependent antigens ^15^. These mice also developed signs of autoimmunity and lymphoproliferation upon aging ^15^. However, as MARK2 is ubiquitously expressed in hematopoietic and non-hematopoietic cells, it was unclear whether the immune phenotypes observed were T cell–intrinsic or result from indirect effects mediated by other cell types. Moreover, in this model, CD8⁺ T cells were not investigated in detail ^15^. This is particularly relevant since MARK2 is more expressed in CD8 than in CD4 T cells and therefore the phenotype of MARK2 deficiency is expected to be stronger in the CD8 compartment. In our study, by using a T cell–specific conditional knockout model (cKO), we addressed these limitations and provided direct evidence that MARK2 plays a T cell–intrinsic role in maintaining immune homeostasis and in controlling sustained T cell activation. We also show that MARK2 deficiency in T cells promotes the development of central memory (cM) CD8⁺ T cells and alters the expression of key surface markers such as CD44 and CD62L on naïve T cells. Bone marrow chimeras experiments confirmed that these phenotypic modifications were a direct consequence of the loss of MARK2 in T cells. Thus, our cKO approach allowed us to study T cell-intrinsic functions of MARK2 that could not be obtained previously. We provide strong evidence that MARK2 functions as a gatekeeper of peripheral T cell homeostasis and protects against the development of autoimmunity. Indeed, MARK2 deficiency in T cells promotes spontaneous autoimmunity in mice upon aging, with high levels of anti-nuclear antibodies (ANA) and lymphocytic infiltration in non-lymphoid tissues. Of note, cohorts of mice of both genotypes were kept for more than 24 months in EOPS conditions and mice never showed obvious health issues or premature death, suggesting that autoimmunity develops gradually. Since we did not detect changes in Treg frequency and phenotype, the breakdown of tolerance likely reflects a defect in conventional T cell-intrinsic mechanisms.

Although we showed that early TCR signaling was not affected by MARK2 deficiency, we observed sustained activation, proliferation, and cytokine production in MARK2^KO^ CD8 T cells, occurring independently of CD28 engagement. This is striking given that CD28 is considered essential for full naïve T cell activation and effector differentiation. In the absence of MARK2, we observed increased expression of CD80 and CD86, the ligands for CD28. Our findings are consistent with recent publications showing that T cells express CD80 and CD86 upon activation and engage CD28 *in cis,* promoting T cell-autonomous co-stimulation. However, the inability of CTLA-4-Ig to block sustained activation in MARK2-deficient T cells suggests that MARK2 neither regulates *cis-*nor *trans-*CD28 signaling. Instead, our data support a model in which MARK2 prevents ligand-independent activation of CD28 signaling.

To better understand the role of MARK2 in T cell biology, we performed single-cell RNA-sequencing (scRNA-seq) on resting or activated naïve CD8 T cells from WT or MARK2^cKO^ mice. Recent studies have shown that naïve T cells are transcriptionally heterogeneous in mice, and this heterogeneity may shape their fate prior to antigen recognition ^29,30^. We therefore chose scRNA-seq to visualize sample heterogeneity at a single-cell level. This high-resolution approach revealed that MARK2-deficient naïve CD8 T cells were transcriptionally different from WT cells, as shown by striking differences in the naïve-like clusters at basal state. MARK2^KO^ naïve T cells show an overrepresentation of a cluster of cells characterized by an enrichment of transcripts involved in the CD28 signaling (*Pik3r1, Pik3cd, Prkcq, Card11*) and glycolysis (*Ldha, Pfkl, Pfkfb3, Eno1*) pathways. This correlates with our observations showing that, in response to TCR activation alone, MARK2-deficient T cells exhibit increased activation of the PI3K-AKT-mTORC1 axis, enhanced glycolysis, proliferation, survival and cytokine production, all responses that are typically induced by CD28 co-engagement in WT cells. Thus, our data suggest that in the absence of MARK2, CD8 T cells are primed to activate a CD28-associated program and that MARK2 is a gatekeeper of the PI3K–AKT–mTORC1 axis. The deregulation of this axis could be the cause of the autoimmunity observed in the MARK2^cKO^ mice. Indeed, defects in genes related to the PI3K-AKT-mTOR pathway are known to cause systemic inflammation and autoimmunity, both in humans and in mice ^31–33^. Further work will be necessary to decipher the precise molecular pathways by which MARK2 prevents *in vivo* immune dysregulation over time.

One of the consequences of the deregulation of the PI3K–AKT–mTORC1 pathway, is the metabolic reprogramming of MARK2-deficient T cells. Since this reprogramming is a key process of T cell fate and function, changes in the baseline metabolic activity of naïve T cells in MARK2^cKO^ mice could contribute to the expansion of central memory cells ^34^. By limiting the activation of the PI3K–AKT–mTORC1 pathway upon TCR triggering, MARK2 prevents T cells from maturing into memory or potentially harmful self-reactive cells in the absence of danger signals.

Although the phosphorylation sequence of MARK2 on serine and threonine is well known ^7,35^, its downstream targets in T cells remain poorly characterized. In pancreatic cells, MARK2 has been shown to phosphorylate and thus, to inhibit CRTC2 nuclear translocation and the subsequent coactivation of CREB ^24^. In this study, we show that, upon TCR stimulation, MARK2 specifically phosphorylates CRTC2 on Ser275, and that this phosphorylation is inhibited by CD28 costimulation. Of note, pharmacological inhibition of CREB also diminishes mTOR activity suggesting a regulatory mechanism between mTOR and CREB. Moreover, MARK2 inactivation results in the upregulation of CREB transcriptional activity, suggesting that MARK2 negatively regulates CREB-induced gene expression through CRTC2 phosphorylation.

CREB has been involved in a variety of cellular processes, including cell proliferation, survival and differentiation, which could support the phenotype observed in MARK2-deficient T cells ^36^. Indeed, our data reveal a novel regulatory pathway by which MARK2 inhibits CREB activation and mTOR signaling in T cells. They suggest that co-stimulation by CD28 relieves this brake by disrupting CRTC2 phosphorylation and CREB function (see schematic representation). This is consistent with a previous study reporting that CREB activation is enhanced by CD3/CD28 signaling pathway ^37^. Yet, other MARK2 substrates may contribute to its function in T cells. For example, HDAC7, a histone deacetylase involved in transcriptional repression, has been shown to be phosphorylated by MARK2 in non-lymphoid cells^38^. Interestingly, in thymocytes, many HDAC7-regulated genes are also components of the CD28 signaling pathway ^39^. Therefore, the phenotype found in our study may result from MARK2 kinase activity on multiple substrates. Further work will be necessary to clarify the role of MARK2 molecular targets in T cells both *in vivo* and *in vitro*.

Our results suggest a paradigm shift in CD28 signaling, whereby CD28 would not only induce a costimulatory signal but also lift a MARK2-dependent inhibitory signal, which precludes sustained activation by the TCR. Of note, the CD28 signaling pathway has already been shown to be negatively regulated. Among others, the E3 ubiquitin ligase CBL-B, which is a negative regulator of T cell activation acts downstream of both TCR and CD28 by targeting signaling proteins such as PI3K and PLCγ1 for ubiquitination and degradation ^40^. Like MARK2, CBL-B deletion induces autoimmunity, lowers the requirement for CD28 co-stimulation and enhances TCR-induced T cell responses ^41^. This strict control of the costimulatory pathway may ensure that TCR signaling is not sufficient to induce full T cell activation in the absence of a “danger signal” and thus protects from autoimmunity.

The prediction of our working hypothesis is that MARK2 is activated by the TCR, which is suggested by several studies ^42–44^, and inhibited by CD28 as suggested by our data showing the decrease of CRTC2 phosphorylation. Investigating MARK2 modifications in response to these two signals will be important to better characterize MARK2 function.

In summary, we identified MARK2 as a key intracellular checkpoint controlling T cell activation, metabolism, and peripheral tolerance. Interestingly, we confirmed several of our findings in human T cells, including enhanced proliferation (this study) and cytokine production ^5^. These findings may have important implications for cancer immunotherapy, particularly in the context of the tumor microenvironment (TME), where T cells can be metabolically suppressed and functionally exhausted. Within the TME, escape mechanisms frequently involve downregulation of CD80 and CD86 on dendritic cells and/or upregulation of inhibitory receptors such as CTLA-4, PD-1, and Tim-3 on T cells, preventing full activation and effector differentiation ^45^. Targeting MARK2 activity could enhance CD28-independent T cell responses in tumors by increasing mTOR dependent-metabolism and effector functions of tumor-infiltrating T cells. Notably, other negative regulators such as CBL-B and HPK1 have already emerged as promising targets to enhance T cell responses in cancer ^46,47^. Further studies will be needed to explore this potential and identify the most effective ways to modulate MARK2 or its downstream pathways in a therapeutic context.

## Methods

### Mice

Animal care and use for this study were performed in accordance with the recommendations of the European Community (2010/63/UE) for the care and use of laboratory animals. Experimental procedures were specifically approved by the ethics committee of the Institut Curie CEEA-IC #118 (Authorization APAFiS#50092-2024060410598283-v1 given by National Authority) in compliance with the international guidelines. Animals were housed under specific and opportunistic pathogen-free (SOPF) conditions and were kept under ambient temperature (21–22 °C) and 50–60% humidity, with 12–12 h on-off light cycle. Both male and female between 6 and 14 weeks of age were used in this study unless specified. For ANA and autoimmunity detection mice between 5 and 24 months were analyzed. All mice were euthanized using cervical dislocation. CD4-Cre and OT-I TCR transgenic mice have been described elsewhere ^17,48^. Rag2-deficient and C57Bl6/J CD45.1 and CD45.2 mice were obtained from Charles Rivers Laboratories.

### Generating Mark2-deficient mice

Mark2-flox mice were generated by the CIPHE consortium. A self-excising ACN cassette was introduced in the intron located between *Mark2* exons 3 and 4, and a cassette permitting the expression of a diphtheria toxin fragment was abutted to the targeting construct. JM8.F6 B6N ES cells were electroporated with the targeting vector. After selection in G418, ES cell clones were screened for proper homologous recombination by Southern blot and PCR analysis. A probe specific for the NeoR cassette was further used to ensure that adventitious nonhomologous recombination events had not occurred in the selected clones. Mutant ES cells were injected into Balb/c blastocysts. The resulting mutant mice are denoted as Mark2-flox mice (B6-Mark2tm1Ciphe). Genotyping of the *Mark2*-flox allele was performed by PCR using two pairs of primers. The first pair (sense 5′-GTAGCTGTGAAGATCATCGAC-3′ and antisense 5′-CTTGAAGGGACTGTTGAGTC-3′) amplified a 138-bp band in the case of the WT allele, whereas we amplified a 233-bp band in the case of the *Mark2*-flox allele. Founders were backcrossed with C57BL/6J mice to remove potential off-targets and to segregate the mutated allele. Homozygous Mark2^flox/flox^ mice were crossed onto CD4-Cre transgenic animals (*CD4-Cre* JAX stock #017336) to obtain Mark2^cKO^ mice. The presence of the Cre transgene was tested by PCR using the following primers: 5’-CGAGTGATGAGGTTCGCAAG-3’ and 5’-TGAGTGAACGAACCTGGTCG-3’ and amplify a 390-bp band if present. When genotyping the CD4-Cre, a 250-bp band was amplified as an internal control using primers specific for the myogenin gene (5’-TGGGCTGGGTGTTAGCCTTA-3’ and 5’-TTACGTCCATCGTGGACAGC-3’).

### Antibodies and reagents

All the antibodies and reagents used in this study are listed in extended table 1.

### Cell preparation and in vitro activation assays

Single-cell suspensions of splenocytes or lymph node cells were obtained by mechanical disruption through a 40µm nylon cell strainer (Fisher Scientific, catalog no. 11587522). Red blood cells were lysed using a hypotonic buffer (Sigma-Aldrich, catalog no. R7757-100ML) for 5 min at room temperature (RT). Naïve CD8 T cells were purified with a mouse Naïve CD8a^+^ T cell Isolation Kit (Miltenyi Biotec, catalog no. 130-096-543), B cells were purified with a mouse B Cell Isolation Kit (Miltenyi Biotech, catalog no. 130-090-862), and CD4 T cells were purified with a mouse CD4^+^ T Cell Isolation Kit (Miltenyi Biotech, catalog no. 130-104-454).

For in vitro assays, cells were cultivated in complete RPMI medium (complete RPMI): RPMI 1640 medium GlutaMAX (Gibco, catalog no. 61870044) containing 10% Fetal Calf Serum (FCS), 1X non-essential amino acids (Gibco, catalog no. 11140035), 1 mM sodium pyruvate (Gibco, catalog no. 11360039), 100 U ml^-1^ Penicillin/Streptomycin (Gibco, catalog no. 15140122), 55 μM 2-mercaptoethanol (Gibco, catalog no. 31350010) and were cultured at 37°C and 5% CO_2_.

For overnight activation analyses, purified naïve CD8^+^ T cells were stimulated with increased doses of G4 peptide (OVA-G4 Peptide, SIIGFEKL, OVA (257-264) Variant, Anaspec, catalog no. 089AS-64384) or N4 peptide (OVA (257-264), AnaSpec, catalog no. 089AS-60193-1), as indicated on the figure. The following day, upregulation of CD69 and CD25 were analyzed by flow cytometry.

For time-course activation analyses, cells were stimulated with N4 peptide (1 nM) in the absence or presence of anti-CD28 1 µg ml^-1^ (Biolegend, catalog no. 102116, Clone 37.51) at a concentration of 0.5 x 10^6^ cells ml-^1^ in complete RPMI.

For supernatant experiments, purified MARK2 ^KO^ naïve OT-I CD8^+^ T cells were stimulated with N4 peptide, as described above, for 72h at 0.1 x 10^6^ 100 ul^-1^ in complete RPMI and supernatant was harvested and frozen at -20°C. Naïve OT-I CD8^+^ T cells were activated as above and 100 ul of thawed previous supernatant was added to the culture.

For the activation of polyclonal naïve CD8^+^ T cells, 24 well plates were pre-coated with anti-CD3, 5 µg ml^-1^, (Biolegend, catalog no. 100340, Clone 145-2C11), and cells were cultured for 72h at 0.5 x 10^6^ cells ml^-1^ in complete RPMI in the absence or presence of soluble anti-CD28 (1 µg ml^-1^).

When indicated in the figure, purified naïve OT-I CD8^+^ T cells were activated with 10 ng ml^-1^ Phorbol 12-myristate 13-acetate (PMA) (Calbiochem, catalog no. 524400, stock solution 1 mg ml^-1^ in DMSO) and 250 ng ml^-1^ ionomycin (Calbiochem, catalog no. 407950, stock solution 1 mg ml^-1^ in DMSO) in complete RPMI for 72h.

### Proliferation assays and cytokine production

For proliferation assay, cells were purified as above and stained with 5 μM CellTrace Violet dye (Invitrogen, catalog no. C34571, DF 1:1,000) at 10 x 10^6^ cells ml^-1^ in PBS for 20 min at 37°C in the dark. Labeled cells were washed and cultured at 37°C and 5% CO_2_ for 72h in round-bottomed 96 well plates at 0.5 x 10^6^ cells ml^-1^ in complete RPMI in presence of N4 peptide (1 nM), with or without anti-CD28 (1 µg ml^-1^). For CTLA-Ig treated conditions, cells were cultured for 72h in 24 well plate at 0.5 x 10^6^ cells ml^-1^ in complete RPMI and activated with N4 peptide (1 nM), in presence of Recombinant CTLA-4-Ig at 5 µg ml^-1^ (Bio X Cell, catalog no. BX-BE0099-1MG) or Recombinant IgG1 control at 5 µg ml^-1^ (Bio X Cell, catalog no. BX-BE0096-1MG). Cell proliferation was analyzed by flow cytometry and quantified using FlowJo software. The proliferation index, as calculated by FlowJo, corresponds to the total number of cell divisions divided by the number of cells that underwent at least one division.

To evaluate T cell cytokine production, cells were activated for 24h with N4 peptide (1 nM), with or without anti-CD28 (1µg ml^-1^), in complete RPMI. The following day, cells were restimulated under the same conditions for 1h, after which Brefeldin A 5 µg ml^-1^, (Sigma Aldrich, catalog no. B5936-200UL) was added to the culture for 4h. Cells were washed and stained for surface markers then fixed and permeabilized for intracellular staining of IFNγ and TNFα (see flow cytometry section for details).

### Hematopoietic chimera

Bone marrow (BM) was flushed from tibia and femur of MARK2^cKO^ mice or control littermates (Mark2^flox/flox^). Four million BM cells from donor groups were mixed and injected into the retro-orbital vein of irradiated Rag2 KO recipients (4.2 Gys). 8 weeks after reconstitution, mice were sacrificed, and spleen and lymph nodes (LN) were collected for FACS analysis.

### Autoimmunity

Serum was harvested from blood collected from the retro-orbital vein and centrifuged at 650g for 10 minutes at room temperature. ANA detection was performed using HEP-2 slides (Kallestad HEp-2 Slides, Bio-Rad, catalog no. 26101). Slides were first incubated with mouse serum diluted at 1:80 and 1:160 in PBS, followed by incubation with AF488-goat anti-mouse IgG (Invitrogen, catalog no. A11029, Highly Cross-Adsorbed). After washing twice with PBS, slides were rinsed in H2O, mounted with 5 μl of Fluoromount-G and dried in the dark at room temperature overnight before microscope acquisition. Images were acquired using an inverted laser scanning confocal microscope (Leica DMi8, SP8 scanning head unit). Immunofluorescence images shown in Fig.3A represent a single plane. Data were analyzed on Fiji software. The ANA score (0= no staining, 5= high intensity staining) was defined in a blinded manner based on fluorescence intensity.

Samples of pancreas, liver, lungs and kidneys were fixed with 10% neutral buffered formalin and embedded in paraffin wax. 4 µm-thick sections were further stained using a routine hematoxylin-eosin-saffron staining for histological analysis. All samples were evaluated by a trained board-certified pathologist in a double-blind manner, and lesions were systematically recorded. The following 7 findings were further semi-quantified according a 3-level scale (with 0=normal; 1=mild change and 2=severe change): kidney cyst and glomerulopathy, hepatopathy, lymphocytic aggregates in kidney, pancreas, lungs and liver. A total score of histological lesions (0-16) was calculated by summing all scores.

### Drugs and inhibitors

Rapamycin (mTOR inhibitor) (Tocris Bioscience, catalog no. 1292) was used at a final concentration of 20 nM and LY 294002 (PI3K inhibitor) (Sigma-Aldrich, catalog no. 440202) at 10 µM final. Both inhibitors were added in the wells at the same time as N4 peptide (1 nM). CREB inhibitor, 666-15, (Calbiochem, catalog no. 5383410001) was added to the culture of 24h activated cells for 3h at a final concentration of 5 µM in complete RPMI.

### Surface and intracellular staining by flow cytometry

Depending on the experiments, samples were acquired either on a Novocyte flow cytometer (Agilent) or on a VYB MacsQUANT (Miltenyi Biotec). Cells were first stained with Live/Dead Fixable Aqua (Molecular Probes, catalog no. L34957), then incubated with FcR Blocking Reagent (Miltenyi Biotec, catalog no. 130-092-575, DF 1:50) for 25 min at 4°C to prevent non-specific antibody binding. Cells were then stained for surface markers using pre-conjugated primary antibodies diluted in FACS buffer (PBS, 0,5% BSA, 1 mM EDTA) for 30 min at 4°C. The following antibodies were used: CD4 (Biolegend, catalog no. 100528, Clone RM4-5, DF 1: 1,000), CD5 (Biolegend, catalog no. 100608, Clone 53-7.3, DF 1:1,000), CD8 (Biolegend, catalog no. 100712, Clone 53-6.7, DF 1:2000), CD25 (BD Pharmingen, catalog no. 553072, Clone 7D4,DF 1:300), CD28 (Biolegend, catalog no. 122007, Clone E18, DF 1:200) CD44 (Biolegend, catalog no. 103008, Clone IM7, DF 1:2,000), CD45.1, (Biolegend, catalog no.110731, clone A20, DF 1:200), CD45.2 (Biolegend, catalog no. 109823, Clone 104, DF 1: 200), CD62L (Biolegend, catalog no. 104418, Clone MEL-14, DF 1:2,000), CD69 (Biolegend, catalog no. 104527, Clone H1.2F3, DF 1:400), CD80 (BD Pharmingen, catalog no. 561955, Clone 16-10A1, DF 1:200), CD86 (BD Pharmingen, catalog no. 561964, Clone GL1, DF 1:200), CD278 (ICOS) (Biolegend, catalog no. 117421, Clone 7E.17G9, DF 1:400), CD279 (PD-1) (Biolegend, catalog no. 135216, Clone 29F.1A12, DF 1:400), CD357 (GITR) (Biolegend, catalog no. 126317, Clone DTA-1, DF 1:2,000), TCRβ (Biolegend, catalog no. 553171, Clone H57-597, DF 1:1,000), TCRVa2 (Biolegend, catalog no. 127825, Clone B20.1, DF 1:400), TCRVβ 5.1, 5.2 (BD Pharmingen, catalog no. 562086, Clone MR9-4, DF 1:500).

For intracellular Foxp3 staining, cells were first stained for surface markers, then resuspended in fixation/permeabilization buffer (Foxp3/Transcription Factor Buffer set, eBioscience-Invitrogen, catalog no. 00-5523-00) for 30 min at 4°C. After washing the cells with permeabilization buffer, staining with PE-anti-Foxp3 (eBioscience-Invitrogen, catalog no. 12-5773-82, Clone FJK-16s, DF 1:100) was performed for 30 min at 4°C.

Intracellular staining of cytokines was conducted using the Cytofix/Cytoperm kit (BD Biosciences, catalog no. 554714), according to the manufacturer’s protocol. Cells were stained with BV421-anti-IFNγ (Biolegend, catalog no. 505829, Clone XMG1.2, DF 1:100) and BV-421-anti-TNFα (Biolegend, catalog no. 506327, Clone MP6-XT22, DF 1:100) for 30 min at 4°C.

For phosflow experiments, cells were harvested, washed once in PBS and immediately fixed for 15 min at 37°C with 4% paraformaldehyde (PFA, ELECTRON MICROSCOPY SCIENCES, catalog no. 15710) to maintain their phosphorylation state. Cells were washed in FACS buffer and permeabilized in BD Phosflow™ Perm Buffer III (BD Biosciences, catalog no. 558050) for 30 min at 4°C. After washing, intracellular staining was performed for 45min at 4°C with AF488-anti-S6 (pS235/236) (Cell signaling, catalog no. 4803S, Clone D57.2.2E, DF 1:20), PE-Cy7-anti-NFκB p65 (pS529) (BD Biosciences, catalog no. 560335, Clone K10-895.12.50, DF 1:20) and unconjugated anti-AKT (pT308) (Cell Signaling, catalog no. 13038T, Clone D25E6, DF 1:100), all diluted in FACS buffer. To detect AKT phosphorylation on T308 by flow cytometry, cells were additionally stained with a secondary AF488-anti-rabbit antibody (Invitrogen, catalog no. A11034, DF 1:200) for 45 min at 4°C in FACS buffer, then washed prior to analysis.

### OVA-tetramer stimulation

The biotinylated monomer H-2Kb-SIINFEKL was produced by the Recombinant protein facility P2R (UMS 016 BioCore, Nantes Biotech, France). For tetramer preparation, streptavidin-PE (Biolegend, catalog no. 405203) was added sequentially to the biotinylated H-2Kb-SIINFEKL monomer solution. For short-term stimulation assays, naïve OT-I CD8^+^ T cells were stimulated with 40 nM of tetramers for 5, 15 or 30 min at 37°C in RPMI. Cells were then washed once in PBS and resuspended in RIPA lysis buffer for Western blot analysis (see the following section for details).

### Western-blot

Unstimulated or activated naïve OT-I CD8^+^ T cells were washed in PBS and resuspended at 20 x 10^6^ cells ml^-1^ in RIPA lysis buffer (Pierce® RIPA Buffer, Thermo Scientific, catalog no. 89901) supplemented with phosphatase inhibitor (Halt™ Phosphatase Inhibitor Cocktail 100X, Thermo Scientific, catalog no. 78426) and protease inhibitors 1X (cOmplete, EDTA-free Protease Inhibitor Cocktail Tablets, Roche, catalog no. 11873580001) for 15 min on ice and treated with nuclease (Pierce Universal Nuclease For Cell Lysis, Thermo Scientific, catalog n°. 88701). Lysates were centrifuged for 12 min at 4°C and supernatants were transferred to new tubes. After adding 4x Laemmli Sample Buffer (Bio-Rad, catalog no. 1610747) and DTT (Pierce DTT No-Weigh (TM) Format, Thermo Scientific, catalog no. A39255), supernatants were boiled for 5 min at 95°C. Samples were loaded on 4–15% Mini-PROTEAN™ TGX Stain-Free™ Protein Gel (Bio-Rad, catalog no. 4568086) and transferred to PVDF membranes (Trans-Blot Turbo RTA Transfer Mini PVDF, Bio-Rad, catalog no. 1704272) with Trans-Blot Turbo Transfer System (Bio-Rad). Membranes were then blocked with TBS, 0,05% Tween-20, 5% BSA for 1h at RT on rocking platform shaker and incubated overnight at 4°C with primary antibodies. The following day, membranes were washed 3 times with TBS, 0,05% Tween-20 and incubated 45 min at RT with peroxidase-conjugated secondary antibodies (Peroxidase Goat Anti-Mouse IgG (H+L), catalog no. 115-035-146, or Peroxidase Goat Anti-Rabbit IgG (H+L), catalog no. 111-035-144, both from Jackson ImmunoResearch Laboratories INC., dilution 1:10,000). Proteins were revealed with Clarity Western ECL substrate (Bio-Rad, catalog no. 1705061), according to the manufacturer’s protocol, and chemiluminescence was detected with a Chemidoc MP imaging system (Bio-Rad). Western blots were quantified with the Image lab software (Bio-Rad).

### Seahorse analysis

All reagents for Seahorse analysis (Seahorse XF Cell Mito Stress Test Kit, catalog no. 103015-100 and Seahorse XF Glycolysis Stress Test Kit, catalog no. 103020-100) were purchased from Agilent and data were acquired using a Seahorse XFe96 bioanalyzer. Purified naïve OT-I CD8+ T cells were stimulated with N4 peptide (1 nM) in the absence or presence of anti-CD28 (1 µg ml^-1^) in complete RPMI at 37°C and 5% CO_2._ After 24h of stimulation, cells were washed in pre-warmed Seahorse XF RPMI medium pH 7.4 supplemented with 10 mM glucose, 1 mM pyruvate and 2 mM glutamine and plated at 0.15 X 10^6^ cells 30ul^-1^ per well in Seahorse poly-D-Lysine (PDL) coated microplates. Cells were briefly centrifuged to promote adhesion to the culture plate, and pre-incubated at 37°C in absence of CO_2_ for 30-45 min. The glycolytic rate of the cells was measured at baseline and in response to Rotenone/Antimycin A (0.5 µM concentration final) and to 2-deoxy-D-glucose (2-DG, 50 mM concentration final). The glycolytic Proton Efflux Rate (glycoPER), as calculated by the Seahorse Wave software, specifically quantifies extracellular acidification resulting from glycolysis by subtracting the mitochondrial contribution to total proton efflux rate.

### SCENITH analysis

The assay was performed as described in Arguello R.J. et al., Cell Metab, 2020 ^23^. SCENITH™ kit 488 containing all reagents were obtained from GammaOmics (https://www.scenith.com/try-it) and used according to the manufacturer’s protocol. Unstimulated purified naïve OT-I CD8^+^ T cells were resuspended in Eppendorf tubes at 5 x 10^6^ cells ml^-1^ in complete RPMI and treated with puromycin and either control, oligomycin, 2-DG, oligomycin+2-DG or Harringtonine for 30 min-1h at 37°C and 5% CO_2_. Cells were then washed, stained for surface markers Vα2, CD8, CD62L and CD44 and fixed/permeabilized using Foxp3 transcription factor buffer set. Puromycin expression was detected using an AF488-anti-puromycin diluted in the permeabilization buffer. Flow cytometry analysis was performed with a VYB MacsQUANT (Miltenyi Biotec). Mitochondrial dependance and glycolytic capacity were calculated as previously described ^23^.

### CREB-Luciferase assay

Purified naïve OT-I CD8 T cells were activated overnight with N4 peptide (1 nM) in complete RPMI at a concentration of 1 x 10^6^ cells ml^-1.^ The following day, 5 x 10^6^ activated cells were co-transfected with 10 µg of pNL[NlucP/CRE/Hygro] plasmid (Promega, catalog no. CS186804) and 3 µg of pGL4.13[luc2/SV40] (Promega, catalog no. E6681) plasmid using the Neon^TM^ electroporator device (Invitrogen) with the following settings: 1900V, 10ms, 2 pulses. Immediately after electroporation, 0.3 x 10^6^ cells were plated per well in pre-warmed complete RPMI without phenol red (Gibco, catalog no. 11835030) in Nunc™ MicroWell™ 96-Well (Thermo Scientific, catalog no. 137101). N4 peptide at 1 nM and CD28 antibody, 1 µg ml^-1^ were added for restimulation. Luciferase activity was measured 24h after transfection using the Nano-Glo Dual-luciferase reporter assay system (Promega, catalog no. N1610) on a Clariostar plate reader (BMG Labtech) according to the manufacturer’s instructions.

### Generation of MARK2 KO primary T cells

Buffy coats from healthy donors were obtained from Établissement Français du Sang following INSERM ethical guidelines. Peripheral blood mononuclear cells (PBMCs) were isolated by Ficoll density gradient centrifugation and CD3^+^ T cells were purified with a Pan T Cell Isolation Kit human (Miltenyi Biotec, catalog no. 130-096-535). CD3^+^ T cells were activated for 3 days with plate-bound anti-CD3, 2.5 μg ml^-1^, (Biolegend, catalog no. 317326, clone OKT3) and soluble anti-CD28, 1 µg ml^-1^ (Biolegend, catalog no. 302934, Clone CD28.2) in complete RPMI. MARK2 inactivation was performed by transfecting 2 x 10^6^ activated cells with a mix of Cas9 (IDT, catalog no. 1081061, stock concentration: 61 pmol ul^-1^) and sgRNA targeting MARK2 (Hs.Cas9.MARK2.1.AC, IDT, stock concentration 50 pmol ul^-1^) with P3 Primary Cell 4D-NucleofectorTM (Lonza, catalog no. V4XP-3032) using the AMAXA 4D-Nucleofector (program EO-115). Transfection was performed using a 1:3 ratio Cas9 to sgRNA. Transfected cells were cultured in complete RPMI supplemented with 500 U ml^-1^ of IL-2 for 3 days and editing efficiency was tested at day 4 post-transfection by western blot using an anti-MARK2 antibody (Cell Signaling, catalog no. 9118, DF 1:1,000). A scramble sgRNA (IDT) was used as a control.

To test the functional effects of MARK2 inactivation in primary T cells, T cells were stained with Cell Trace Violet (Invitrogen, catalog no. C34571, DF 1:1,000) and restimulated for 3 days with coated anti-CD3 2.5 µg ml^-1^ or coated anti-CD3 plus soluble anti-CD28 1 µg ml^-1^, in flat bottom 96 well plate. Cell proliferation was analyzed by flow cytometry for CTV dilution.

### Sample preparation for single-cell RNA sequencing (scRNA seq)

Single-cell suspensions of splenocytes from MARK2^cKO^ or WT mice (n=1) were enriched for total CD8^+^ T cells using a mouse CD8^+^ T cell Isolation Kit (Miltenyi Biotec, catalog no. 130-104-075). Cells were then stained, and naïve CD8+ T cells were sorted based on the high surface expression of TCRβ, CD8, CD62L and on the low expression of CD44 using a flow cytometer BD FACSAria (BD Biosciences). Purified naïve CD8^+^ T cells were left unstimulated (T0) or stimulated in 24 well plates coated with anti-CD3 (5 µg ml^-1^) at 0.5 x 10^6^ cells ml^-1^ in complete RPMI for 5hours (T5) or 24hours (T24) at 37°C and 5% CO_2_. Approximately 10,000 cells per condition were loaded onto the 10X Genomics Chromium Controller (Chip G). ScRNA seq librairies were generated using the Chromium Single Cell 3’ Reagent Kit v3.1 (10X Genomics), according to the manufacturer’s instructions (user guide CG000315 Rev C).

### Single-cell RNA sequencing analysis

Sequencing was performed by the Next-Generation Sequencing (NGS) core at the Institut Curie on a NovaSeq6000 Illumina platform. CellRanger v7.1.0 (10X Genomics) software was used to perform alignment, filtering, barcode counting, and Unique Molecular Identifier (UMI) counting. Reads were aligned to the mouse mm10 genome. Datasets were analyzed using RStudio (version 2024.12.0+467) and Seurat package ^49^ (version 5.2.1). For unstimulated and 5h-stimulated conditions, we filtered out cells with < 1,500 and > 6,000 detected genes and cells with < 1,500 and > 25,000 UMI counts. For 24h-stimulated conditions, we filtered out cells with < 1,500 and > 9,000 detected genes and cells with < 1,500 and > 100,000 UMI counts. For all conditions, cells with more than 5% mitochondrial gene expression, and cells with < 15% and > 50% ribosomal gene expression were removed from the analysis. All samples were merged on a single Seurat object and normalized with the standard Seurat workflow (NomalizeData, FindVariableFeatures, ScaleData). PCA was run using Seurat’s RunPCA function on the 3,000 most variable genes identified using the “vst” method of FindVariableFeatures function. All samples were integrated together using Harmony ^50^ method with the Seurat’s RunHarmony function. Clustering was performed with Seurat’s FindClusters function, based on the 20 first principal components from PCA dimensionality reduction, with a resolution parameter set to 0.2. The UMAP was run using Seurat’s RunUMAP function on the 20 first principal components from PCA. Markers for each cluster were identified using Seurat’s FindAllMarkers function and filtered based on an adjusted p-value <0.05. Markers distinguishing naïve-like 1 and naïve-like 2 clusters were identified using Seurat’s FindMarkers function and filtered based on an adjusted p-value <0,05 and an average log2FC > 0.25. Pathway analysis was performed using the Reactome pathway repository available on String website (https://string-db.org). Gene signatures were computed manually and analyzed using the packages SignatuR (https://github.com/carmonalab/SignatuR) and UCell ^51^. Data were visualized using Seurat’s ViolinPlot, Ggplot, FeaturePlot, FeaturePlot_scCustom and Plot_Density_Custom functions.

### Statistical analyses

The number of replicates and independent experiments, and statistical tests used are indicated in the figure legends. Statistical analysis was performed with GraphPad Prism software. Normality of the data was determined using the Shapiro-Wilk test. For comparisons between two groups, a t-test was used. Statistical significance was determined by paired or unpaired t-tests for normally distributed data or, by Wilcoxon or Mann-Whitney tests otherwise. Analysis of activation markers over time was performed with two-way ANOVA with Sidak’s correction for multiple testing. In scRNA seq datasets, statistical comparisons of the proportion of cells in the different clusters from WT and KO samples was determined with the R package “ScProportion Test” (permutation test).

## Aknowledgements

We thank S.Lameiras and M.Bohec (ICGex-NGS platform) for technical help with ScRNAseq; G. Gentric for helping with Seahorse technology; C.Sedlick for the N4-monomer; the mouse facility of Institut Curie and C.Thoule, A.Cros and A.Estelle for animal care, as well as the flow cytometry platform of Institut Curie; R.de Almeida and O.Lantz for helpful discussion; and R. Montagne for advices on scRNAseq analysis.

## Author Contributions

LB and CH conceived and supervised the study.

HF performed all the experiments with some help of LB and SD. FM performed the experiments in human T cells. CG helped with the ScRNA seq analysis. AZ, TC, NP helped for the characterization of the role of MARK2. TL and EB performed the histological analysis of autoimmunity in mice. FF. and CIPHE consortium produced the MARK2^flox/flox^ mouse model.

HF, LB and CH wrote the manuscript with the help of SD for the Mat et met section.

---

**Extended data Fig. 1.**
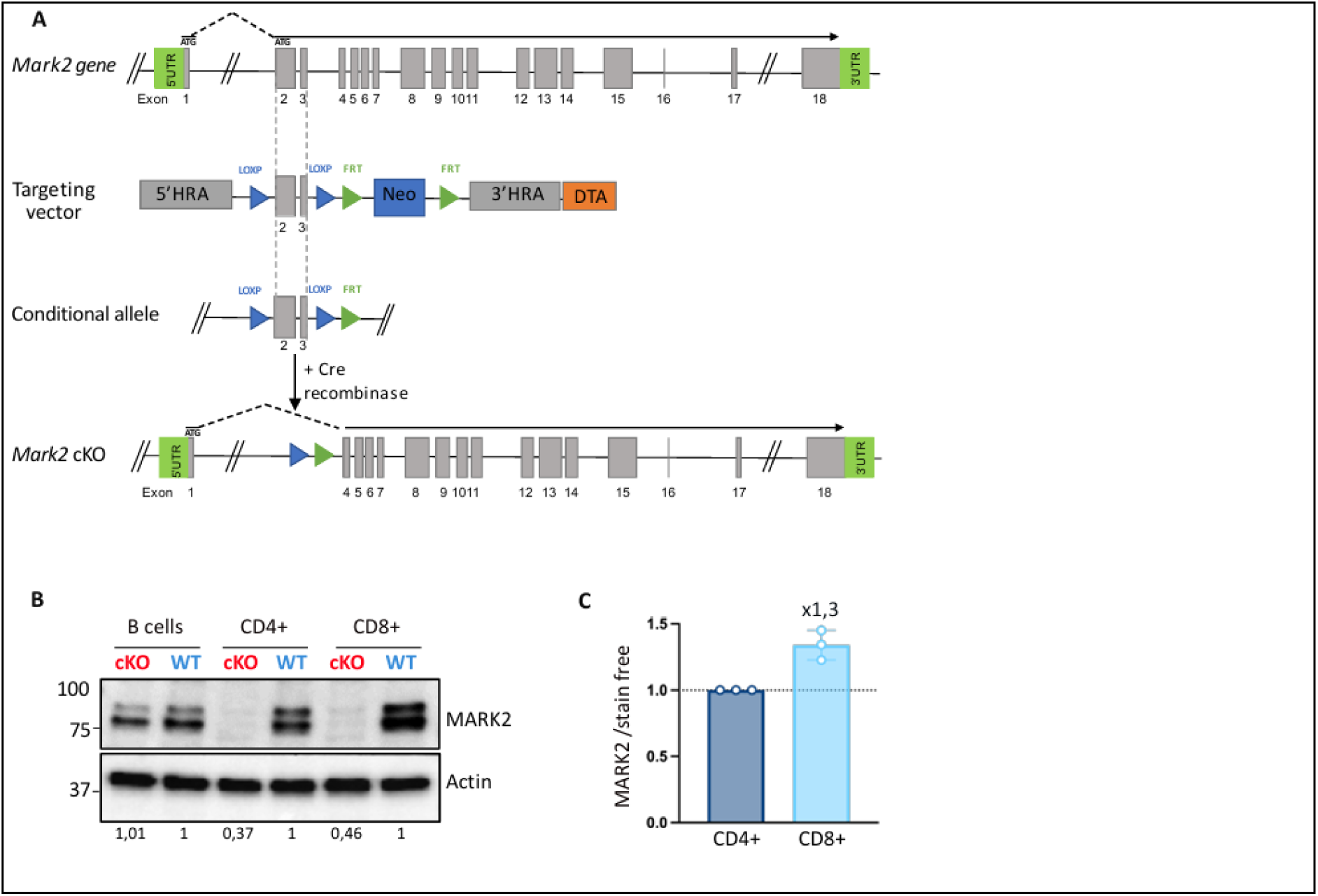
Generation of the MARK2^cKO^ mouse model. **(A)** Schematic representation of the gene targeting strategy used for generating MARK2^cKO^ mice. The targeting vector contained LoxP and FRT sites (blue and green arrowheads, respectively). The neomycin (Neo) and the diphtheria toxin (DTA) cassettes, used for positive and negative selection in ES cells, are shown. Neo cassette was removed after crossing chimeric mice with animals carrying the Flp-e deleter allele. Mice were crossed onto CD4-Cre transgenic mice to achieve the T-cell specific deletion of MARK2. **(B)** Representative immunoblot (n=3) of total protein lysates from B cells, CD4^+^ and CD8^+^ T cells isolated from WT or MARK2^cKO^ mice. Immunodetection was performed using antibodies specific to MARK2 and actin (loading control). Numbers represent the quantification of MARK2 protein in each cell type relative to WT level. **(C)** Quantification of MARK2 band intensity in CD8^+^ T cells relative to CD4^+^ T cells and normalized to total protein levels (stain free).

**Extended data Fig. 2.**
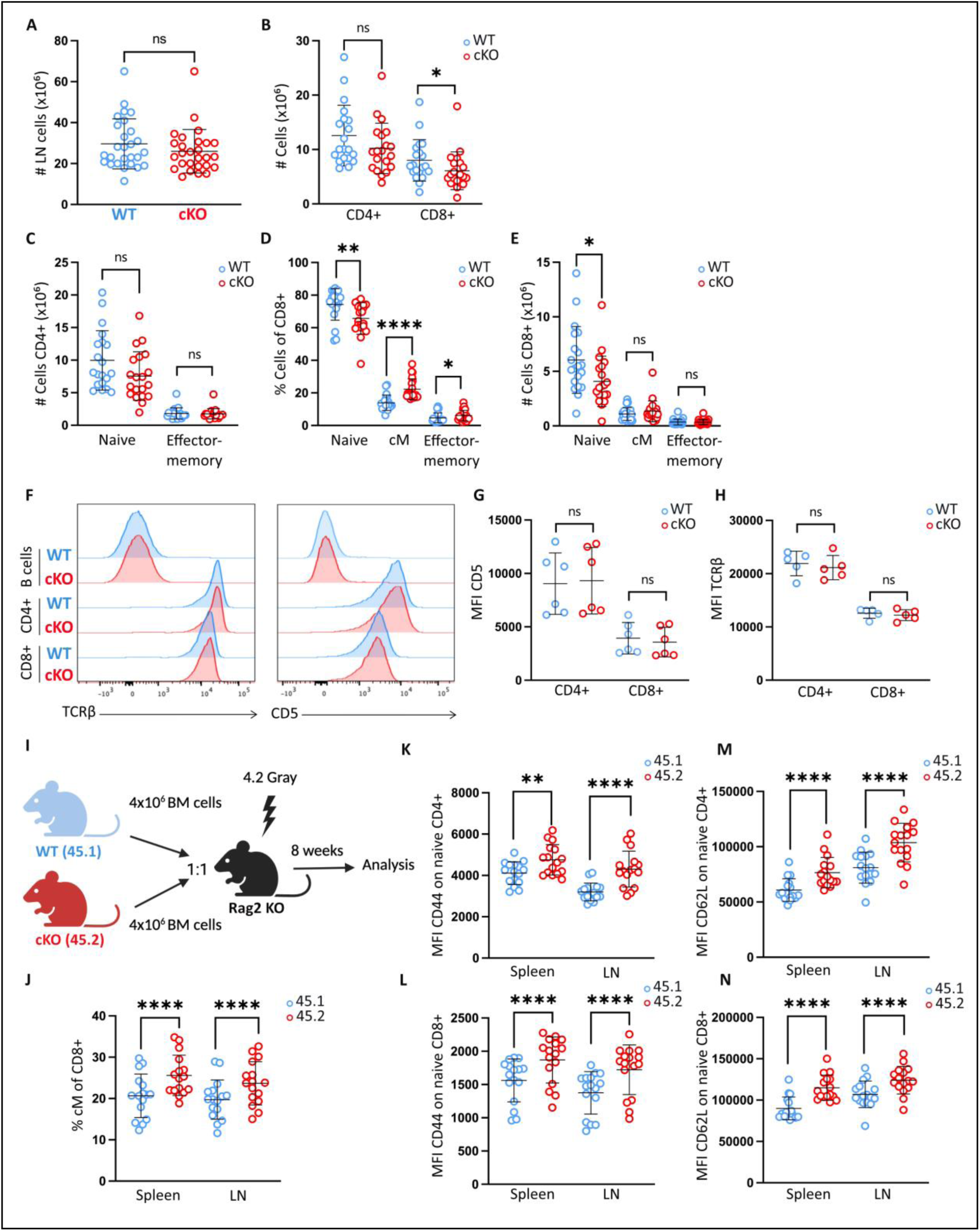
MARK2 controls the formation of central memory (cM) CD8+ T cells in a T-cell intrinsic manner. **(A to E)** Flow cytometry analysis of lymph nodes (LN) harvested from WT (in blue) or MARK2^cKO^ mice (in red)**. (A)** Total LN cell numbers. **(B)** Absolute numbers of CD4^+^ and CD8^+^ T cells. **(C)** Absolute numbers of naïve and effector memory subsets gated on CD4+ T cells**. (D, E)** Frequency and absolute numbers of naïve, cM and effector memory subsets gated on CD8^+^ T cells. **(F to H)** Flow cytometry analysis of the spleens harvested from WT or MARK2^cKO^ mice. **(F)** Representative histograms of TCRβ and CD5 expression on gated B cells, CD4^+^ and CD8^+^ T cells. **(G)** MFI quantification of CD5 and **(H)** TCRβ gated on CD4^+^ and CD8^+^ T cells. **(I to N)** Mixed bone marrow (BM) chimeras mice were generated by injecting equal numbers of WT (45.1) and MARK2^cKO^ (45.2) BM cells into irradiated Rag2^KO^ mice. 8 weeks later, spleens and LN were harvested from reconstituted chimeric mice and were analyzed by flow cytometry. **(I)** Schematic representation of the experiment. **(J)** Percentage of cM cells among CD8+ T cells from chimeric mice. **(K, L)** Quantification of the MFI of CD44 and **(M, N)** CD62L on gated naïve CD4^+^ and CD8^+^ T cells from chimeric mice. Each dot represents an individual mouse (n=20-27 mice in **A-E**, n=5-6 in **F-H,** n=16 in **I-N).** In graphs, bars show mean and SD. Data were generated from at least n=3 independent experiments. For **A-E**, statistical significance was determined by unpaired t-test for normally distributed data or, by Mann-Whitney test otherwise. In **B**, *P=0.0283. In **D**, **P=0.0021, ****P < 0.0001 and *P=0.0422. In **E**, *P=0.0181. For the mixed BM chimeras experiment, statistical significance was determined by paired t-test for normally distributed data or, by Wilcoxon test otherwise. In **K**, **P=0.016. In **J-N**, ****P < 0.0001.

**Extended data Fig. 4.**
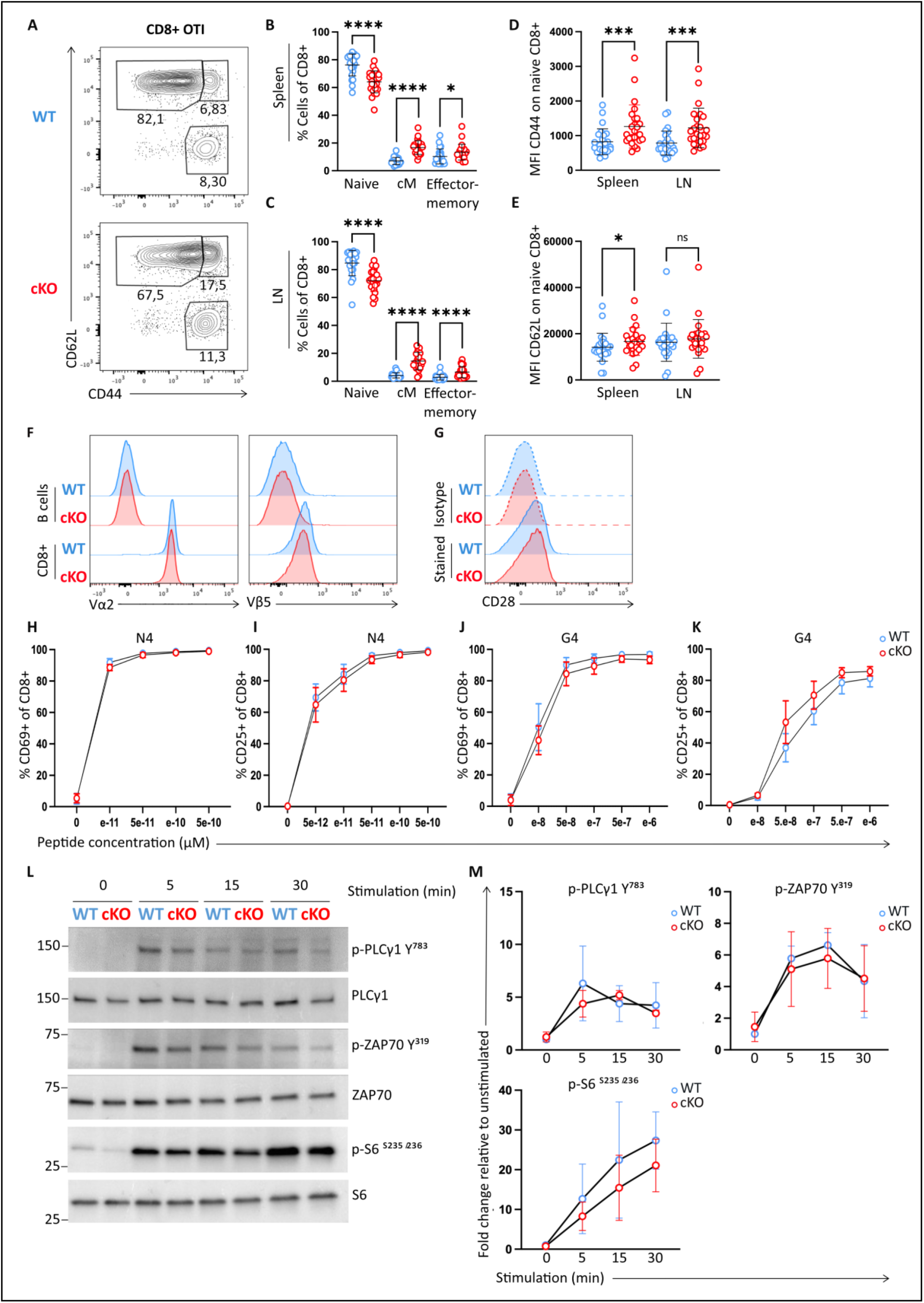
MARK2 does not regulate early T cell responses to antigen-specific stimulation. **(A to G)** Flow cytometry analysis of the spleens and LN harvested from WT or MARK2^cKO^ OT-I mice. **(A)** Representative plots of CD62L and CD44 expression among CD8^+^ T cells isolated from the spleen of OT-I mice. Numbers in quadrant indicate the proportion of the different T cell subsets, described in figure 2C. **(B)** Percentage of naïve, cM and effector memory CD8+ T cells from the spleens or **(C)** the LN of WT or MARK2^cKO^ OT-I mice. **(D)** MFI quantification of CD44 and **(E)** CD62L expression on naïve CD8+ T cells from spleens and LN of WT or MARK2^cKO^ OT-I mice. **(F)** Representative histograms showing expression of TCR Vα2, Vβ5 and **(G)** CD28 on CD8^+^ T cells from WT or MARK2^cKO^ OT-I mice. **(H to M)** Percentages of CD8^+^ OT-I cells expressing CD69 and CD25 upon overnight stimulation with increased concentrations of **(H, I)** SIINFEKL N4 or **(J, K)** SIIGFEKL G4 agonist peptides. **(L)** Representative immunoblot of total protein lysates from WT or MARK2^KO^ naïve CD8+ OT-I T cells stimulated with N4/H-2Kb tetramer (40nM) for 5, 15 or 30 minutes. Immunodetection was performed using antibodies specific for phospho-PLCγ1 (Y^783^), phospho-ZAP70 (Y^319^), phospho-S6 (S^235/236^) and their corresponding total proteins. **(M)** Quantification of phospho-protein band intensities relative to unstimulated WT cells and normalized to total protein levels (stain free). Each dot represents an individual mouse (n=24 mice for **B to E**). Data were generated from at least n=3 independent experiments. Statistical significance was determined by unpaired t-test for normally distributed data or, by Mann-Whitney test otherwise. In **B**, *P=0.0283. In **B and C**, ****P < 0.0001. In **D**, ***P=0.0009 for the spleen and ***P= 0.0005 for the LN. In **E**, *P=0.0371. In **H to M**, graphs show mean and SEM.

**Extended data Fig. 4 bis.**
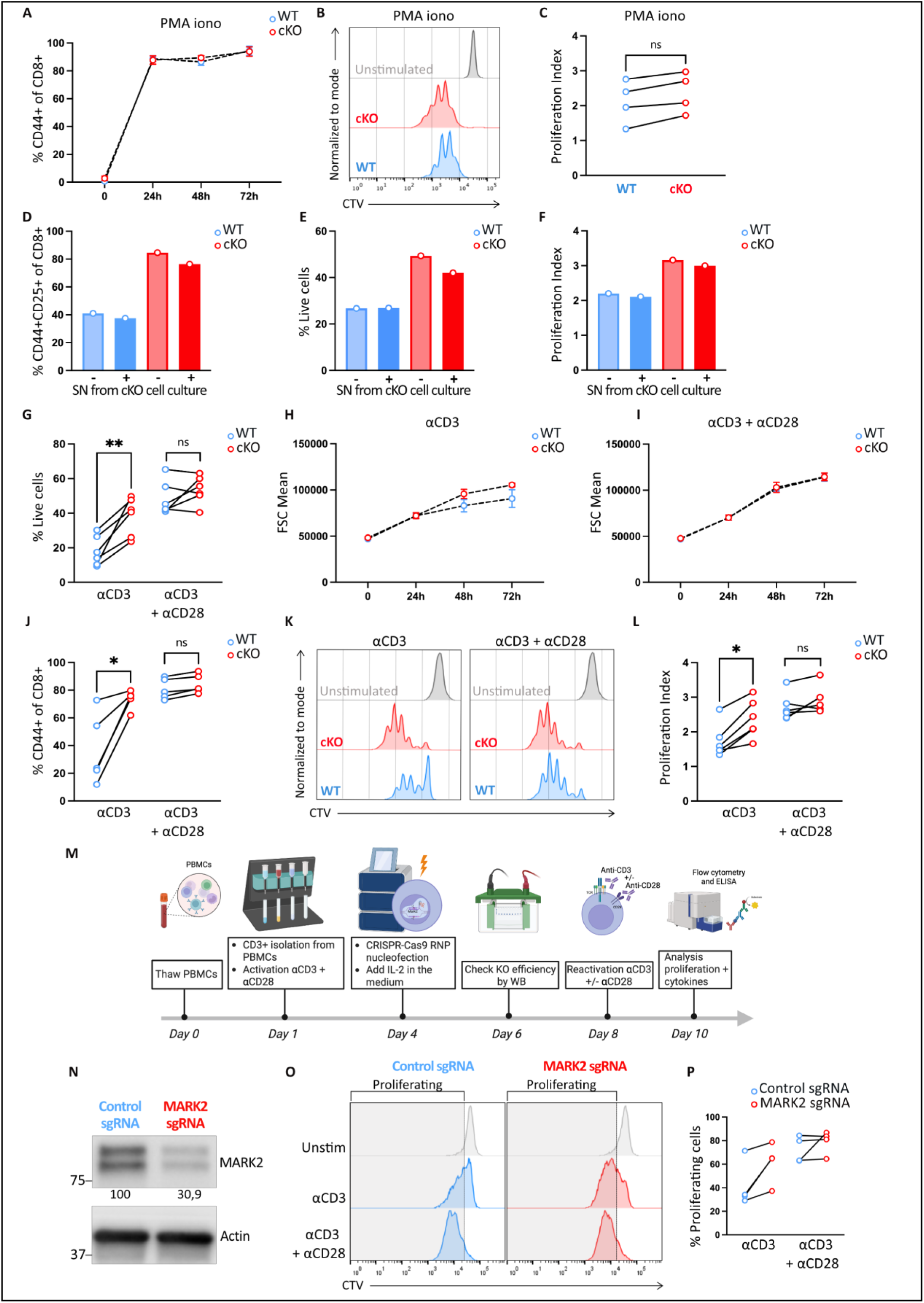
MARK2 ^KO^ CD8^+^ T cells from polyclonal mice and human T cells inactivated for MARK2 by CRISPR-Cas9 acquire a CD28-independent phenotype upon activation. (A to. **C)** Naïve CD8^+^ T cells from WT or MARK2^cKO^ OT-I mice were stimulated *in vitro* for 24, 48 or 72 hours with PMA and ionomycin (PMA-iono). **(A)** CD44 expression and **(B, C)** proliferation index, were assessed by flow cytometry. **(D to F)** WT or MARK2 ^KO^ naïve CD8^+^ T cells were stimulated for 72h with N4 peptide (1nM) and incubated with supernatant from independent cell culture of activated MARK2 ^KO^ T cells. **(D)** Frequency of activated cells expressing CD44 and CD25, **(E)** cell viability, and **(F**) proliferation index, were quantified by flow cytometry. **(G to L)** Flow cytometry analysis of naïve CD8^+^ T cells isolated from WT or MARK2^cKO^ polyclonal mice stimulated *in vitro* for 24, 48 or 72-hours with plate-bound anti-CD3 alone (5μg/ml), or in combination with soluble anti-CD28 (1μg/ml). **(G)** Cell viability measured at 72h following stimulation with anti-CD3 alone or in combination with anti-CD28. **(H)** Cell size over time (as indicated on the x axis) following activation with anti-CD3 alone or **(I)** in presence of anti-CD28. **(J)** CD44 expression and **(K, L)** proliferation index at 72h, were quantified for each condition. **(M to Q)** Human CD3^+^ cells were isolated from PBMCs and were inactivated for MARK2 by CRISPR Cas9. Four days later, cells were reactivated with plate-bound anti-CD3 alone or in presence of anti-CD28 and analyzed by flow cytometry. **(M)** Schematic representation of the CRISPR-Cas9 workflow used to inactivate MARK2 in human T cells. **(N)** Representative immunoblot of total protein from T cells inactivated with control (in blue) or specific MARK2 sgRNA (in red). Immunodetection was performed using antibodies specific for MARK2 and actin (loading control). **(O)** Representative histograms of CTV dilution at 72h post-reactivation. Gray boxes indicate proliferating cells and were quantified in **(P).** Each dot represents an individual mouse (n=4 mice in **A-C**, n=6 in **G-J-L**, n=4 in **H-I**, n=3 in **M-P**). Data were generated from at least n=3 independent experiments, except for **D-F** which is representative of n=2. Statistical significance was determined by unpaired t-test for normally distributed data or, by Mann-Whitney test otherwise. In **G**, **P=0.0051. In **J**, *P=0.0306. In **L**, *P=0.0390.

**Extended data Fig. 6.**
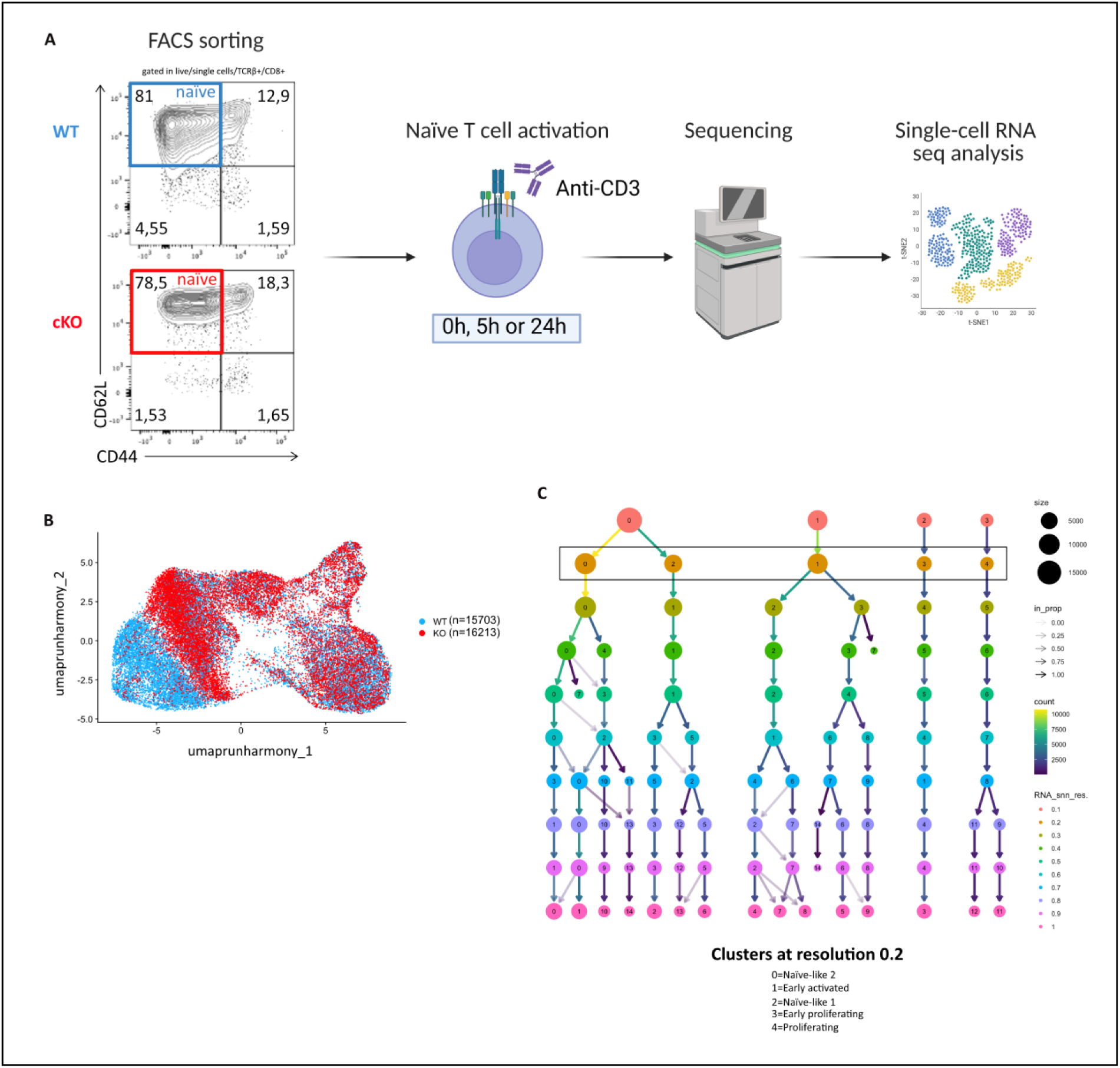
Single-cell RNA sequencing (sc-RNA seq) analysis of resting or activated MARK2 ^KO^ and WT naïve CD8^+^ T cells. **(A)** Schematic representation of the sc-RNA seq workflow from cell purification to data analysis. Naïve CD8+ T cells were sorted based on the high expression of the CD62L marker and on the low expression of CD44. Cells were left unstimulated or activated by the TCR for 5h or 24h before proceeding to the sequencing and analysis. **(B)** UMAP representation showing cell clustering by genotype. **(C)** Clustree representation showing the resolution parameter used for scRNA-seq analysis.

